# Fusion Peptide Insertion Configuration Determines the Mechanism of Membrane Curvature Induction in Human Parainfluenza Virus Type 3

**DOI:** 10.64898/2026.10.07.757304

**Authors:** Melanie A. Brunet, Laura J. S. Lopes, Nina Mansfield, Avijit Sardar, Tara C. Marcink, Paul S. Blank, Myriam Cotten, Anne Moscona, Alexander J. Sodt, Joshua Zimmerberg

**Affiliations:** Eunice Kennedy Shriver National Institute of Child Health and Human Development, National Institutes of Health, Bethesda, Maryland; Center for Host-Pathogen Interaction, Columbia University Vagelos College of Physicians and Surgeons, New York, New York; Department of Biochemistry and Biophysics, Oregon State University, Corvallis, Oregon; Department of Microbiology and Immunology, Department of Physiology and Cellular Biophysics, Columbia University Vagelos College of Physicians and Surgeons, New York, New York; Department of Pediatrics, Columbia University Vagelos College of Physicians and Surgeons, New York, New York

## Abstract

Viral entry requires remodeling the host membrane with a short fusion peptide (FP) that anchors and deforms the target lipid bilayer. However, the contribution of the FP throughout the full fusion pathway remains unknown. Here, all-atom molecular dynamics simulations define the membrane-deformation mechanisms associated with two distinct insertion configurations of membrane-bound human parainfluenza virus type 3 (HPIV3) FP. Molecular simulations quantifying the linear response of curvature to the peptide are complemented with analyses of lipid organization and FP-lipid interactions to define the underlying membrane-deformation mechanisms. The surface-parallel FP configuration generates positive membrane curvature through deformation of the proximal leaflet, consistent with the classical hydrophobic insertion (wedging) mechanism. In contrast, the transmembrane FP configuration generates curvature through localized deformation of the *distal* leaflet, a distinct mechanism that would strongly influence later stages of fusion. Cryo-electron tomography shows that target membrane curvature at virus-host contact sites is consistent with the molecular dynamics prediction. In the absence of F-protein activation and FP insertion, the cryo-ET-determined curvature distribution shifts toward negative values, thereby isolating the role of the fusion protein in inducing early-stage curvature that enables the host and virus membranes to approach one another. Modeling key mutational changes isolates how aromatic and electrostatic interactions act together to promote curvature generation. Taken together, these findings establish FP insertion configuration as a determinant of productive membrane curvature across the full fusion pathway.

## Introduction

Human parainfluenza virus type 3 (HPIV3) is an enveloped paramyxovirus that is a leading cause of lower respiratory tract disease in infants, children, and immunocompromised people (1–3). It enters host cells through membrane fusion mediated by a complex formed by the viral receptor-binding glycoprotein (HN) and fusion glycoprotein (F) (4–6). Upon engagement of a cellular receptor, HN triggers F to undergo a series of structural transitions, exposing the fusion peptide (FP) (7, 8). These conformational changes drive fusion of the viral and cellular membranes, enabling delivery of the viral genome into the host-cell cytoplasm (9, 10). Membrane fusion requires progression through a series of highly curved membrane intermediates (11), including membrane protrusion, hemifusion, and fusion pore formation. (12, 13) Central to this process is the FP, a conserved hydrophobic segment located at the N-terminus of the activated F protein (14) that inserts into and deforms the host membrane (12). However, the orientation and depth adopted by the FP within the membrane is challenging to observe experimentally. Therefore, defining how FP insertion depth and orientation influences membrane curvature generation is essential for understanding viral membrane fusion.

Because membrane curvature provides a mathematical description of membrane shape, understanding how FPs generate and stabilize membrane curvature is critical for defining the physical mechanisms by which viral FPs promote membrane fusion (15, 16). While the molecular mechanism of convex (positive) curvature generation by amphipathic helices (AHs) appears well understood (17, 18), quantifying the sequence- and lipid-dependence of this curvature effect remains a current challenge. Moreover, positive distal leaflet curvature has limited relevance to the entire fusion pathway, which at most stages is favored by concave (negative) leaflet curvature. Characterizing curvature generation by the FP in configurations beyond simple amphipathic, surface-bound configurations is therefore a key target of the modeling portion of this study.

Viral FPs have been reported to adopt distinct membranebound configurations depending on the viral system and membrane environment (19). Differences in insertion configuration suggest that these configurations perturb surrounding lipids differently and thereby influence the mechanisms by which membrane curvature is generated. For surface-parallel peptides, membrane deformation is commonly described by the hydrophobic insertion (wedging) mechanism, which provides a mechanistic basis for peptide-induced curvature generation (17). In this model, insertion of an amphipathic or hydrophobic peptide (20) into a single leaflet generates local packing asymmetry that can drive curvature generation. The sign of the resulting curvature depends on the depth of insertion relative to the leaflet so-called *neutral surface*, the pivotal height at which the leaflet bends (17, 21). In contrast, much less is known about the membrane-deformation properties of FPs in a transmembrane configuration. Experimental studies demonstrated that the parainfluenza virus 5 (PIV5) FP can adopt a transmembrane configuration under specific membrane conditions using IR dichroism (22) and solid-state NMR spectroscopy (23), establishing that a transmembrane FP state is physically feasible. Complementary computational and biophysical studies further characterized oligomeric assemblies of the transmembrane FP and showed that they alter lipid organization, promote water penetration, and alter bilayer properties in ways that the authors suggest will promote membrane fusion between the viral envelope and the host cell membrane(22, 24). These findings demonstrate that transmembrane FP insertion can substantially perturb the host membrane and change bilayer properties, motivating further investigation into how FP orientation influences membrane deformation.

The HPIV3 FP shares a high degree of sequence similarity with the PIV5 FP, suggesting that the two FPs may exhibit related membrane-interaction properties. Over the first 26 residues, the FPs share 76.9% sequence identity (20/26) (22, 24). Both FPs also contain a conserved GXXXG motif near the N-terminus (G3XXXG7), a sequence feature associated with close packing of transmembrane helices and potentially relevant to FP self-association (22). In addition, both FPs exhibit comparable mean hydrophobicity (0.67 and 0.76) and hydrophobic moment (*µH* ≈0.18 − 0.19) and an identical predicted hydrophobic face (VIA AAGAA), features that are consistent with similar membrane insertion behavior. These similarities suggest that a transmembrane insertion state may also be accessible to the HPIV3 FP. Despite these similarities, differences in several residues may alter FP orientation, membrane interactions and lipid perturbation, making HPIV3 a distinct system for characterizing sequence-dependent lipid structure and mechanical stress. Although transmembrane FP configurations have been observed, it remains unclear which membrane-bound orientations are adopted during different stages of the fusion process and whether transitions occur between them. Thus, although prior work on PIV5 provides a rationale for considering a transmembrane insertion configuration in HPIV3 (22, 23), whether multiple FP configurations generate membrane curvature through distinct mechanisms remains unknown.

We propose a model in which the FP initially adopts a surface-parallel configuration (Fig 1 A–B) and subsequently adopts a transmembrane configuration (Fig 1 C–D) during membrane fusion. The physical plausibility of such a transition is supported by recent simulation studies showing that membrane-associated helices can adopt both surface-parallel and transmembrane orientations (25, 26). Here, we focus on the curvature-generating properties of these two insertion configurations rather than the transition between them (25, 26). In the surface-parallel configuration, the FP associates with the proximal leaflet in the membrane, consistent with the behavior of amphipathic helices that preferentially partition into the interfacial region of lipid bilayers (Fig 1 A–B) (27). FP insertion in this configuration (see Fig 1B) is expected to generate curvature through the hydrophobic insertion (wedging) mechanism (17). In the transmembrane configuration, the FP spans the bilayer in a configuration analogous to that observed for PIV5 FP (Fig 1 C–D) (22, 23). The termination of the transmembrane FP directly at the hydrophobic interface on the distal leaflet suggests the possibility that it contributes to a curvature-inducing, lipid-perturbing stress (Fig 1 D). N-terminal phenylalanines and a charged N-terminus at the distal leaflet interface would be key elements of this perturbation.

**Fig. 1.**
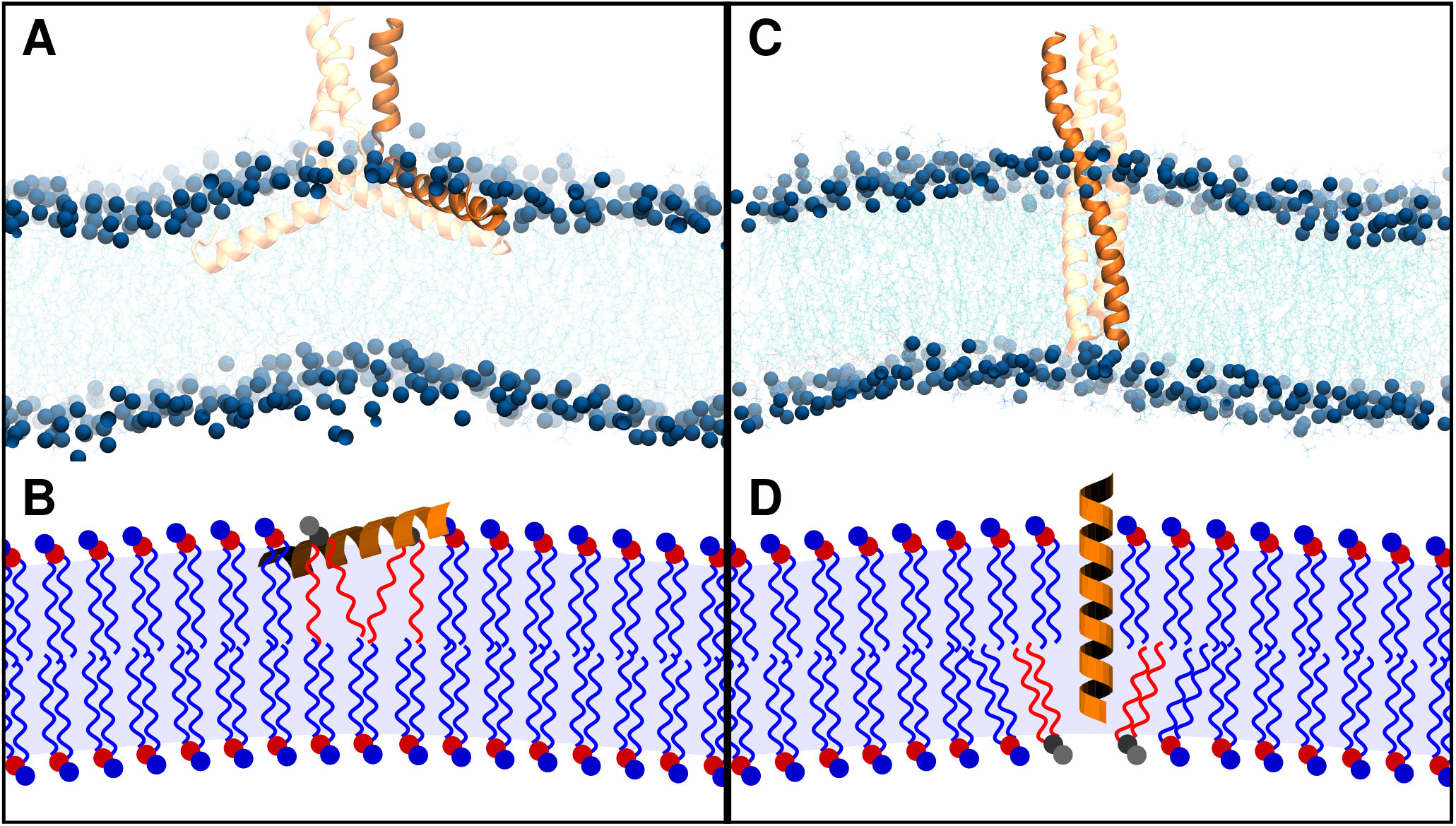
Proposed mechanism of HPIV3 FP-mediated membrane curvature for the surface-parallel and transmembrane configurations. (A) Surface-parallel configuration for the FP trimer. (B) In this configuration, shallow amphipathic insertion above the neutral surface generates positive curvature through localized lipid rearrangement within the proximal leaflet, consistent with the hydrophobic insertion (wedging) mechanism of membrane curvature. (C) Transmembrane configuration for the FP trimer. (D) In this configuration, the FP adopts a transmembrane orientation, generating negative curvature in the distal leaflet, resulting in positive membrane curvature through distal leaflet tail splay and interactions between the inserted N-terminus and surrounding lipids.

Cryo-electron tomography (cryo-ET) can directly visualize and quantify membrane structural changes in systems close to living conditions and provides an experimental complement to precisely controlled molecular simulations. Using fusion inhibitory peptides the HPIV3 fusion process can be stalled after FP insertion (28, 29), yielding geometric information on viral and cellular curvature near the contact site. The pairing of simulation and cryo-ET therefore offers a route to characterizing the influence of the FP and other elements on membrane curvature, if stable intermediates can be frozen while the characteristics of the FP are perturbed. However, the likely dynamic structure of the FP and lipid matrix make direct structural characterization of the FP challenging. Therefore, identifying which elements of the FP sequence influence curvature generation is important for both for understanding how the FP promotes fusion and for inferring its role in the curved intermediates visualized by cryo-ET.

Here, all-atom simulations quantify the curvature induction of the surface-parallel and transmembrane FP configurations, and characterize the lipid structural changes that suggest a molecular mechanisms for curvature generation. Membrane curvature was quantified using the Spatial Extent (SPEX) method (30, 31) in the MembraneAnalysis.jl software package (32, 33). We further analyzed cryo-ET data to compare simulation-predicted membrane curvature with target membrane deformation at virus-host contact sites. Additional simulations with N-terminal modifications to the FP were performed to evaluate the contributions of the N-terminal phenylalanines and the charged N-terminus. This study shows that FP orientation strongly influences membrane curvature, with surface-parallel and transmembrane configurations generating curvature through distinct mechanisms, while membrane composition and specific FP-lipid interactions modulate curvature magnitude. Cryo-ET revealed target membrane deformations consistent with the positive curvature predicted by the simulations, whereas the absence of F-protein activation and FP insertion shifted the curvature distribution toward *negative* values. Altogether, these findings indicate that HPIV3 FP insertion configuration determines the mechanism of membrane curvature generation, while sequence-dependent FP-lipid interactions influence the magnitude of curvature.

## Methods

### System Build and Simulations

All-atom HPIV3 FP-membrane systems were constructed using the CHARMM-GUI Membrane Builder module (34). Membranes containing 100:0, 90:10, and 70:30 mol% POPC:cholesterol were constructed with initial lateral dimensions of approximately 150Å× 150 Å. Both HPIV3 FP monomer and trimer FP systems were generated in either surface-parallel or transmembrane configurations. The monomer FP contained the sequence FFGGVIGTIALGVATSAQITAAVALVEAKQAR and was simulated in the 90:10 and 70:30 mol% POPC:cholesterol membranes. The trimeric FP contained the sequence FFGGVIGTIALGVATSAQITAAVALVEAKQAR SDIEKLKEAIRDTN and was simulated in all three membrane compositions.

The trimeric construct included an extended N-terminal heptad-repeat region (HRN) segment to better represent the structural context of the FP within the intact F protein and to avoid modeling the FP as an isolated peptide. In contrast, the monomer construct contained a shorter HRN segment to avoid introducing a longer isolated HRN region lacking the stabilizing interactions present in the trimer. Systems were solvated with TIP3P water and ionized to final concentrations of 110 mM NaCl and 5 mM KCl. Protonation states were assigned by CHARMM-GUI at pH 7.4. F1A/F2A substitute systems were generated using the CHARMM-GUI mutation option by replacing residues 1 and 2 with alanine (residues F110 and F111 in the native HPIV3 F sequence), and neutral N-terminus (NNT) systems were generated by applying the Neutral N-terminus patch. Three independent replicas were generated for each system.

Molecular dynamics simulations were performed using NAMD 3 (35) with the CHARMM36m (36) force field in the NPT ensemble at 310.15 K and 1 atm. Temperature was maintained using Langevin dynamics and pressure was controlled using the Nosé–Hoover Langevin piston method with semiisotropic pressure coupling. Long-range electrostatic interactions were treated using the particle mesh Ewald method, and van der Waals interactions were force-switched between 10 and 12 Åwith a 12Åcutoff. Bonds involving hydrogen atoms were constrained, allowing a 2 fs integration timestep. Simulation lengths ranged from approximately 2 to 5µs depending on the system. Simulation lengths for each system are listed in Table I.

To maintain the helical structure of the HRN region, harmonic alpha-helical restraints were applied using the collective variables module (37). In monomer systems, restraints were applied to residues 28-32. In trimer systems, restraints were applied independently to residues 35-46 of each protomer. For surface-parallel trimer systems, a harmonic RMSD restraint was applied to the HRN stem region instead of alpha-helical restraints. Restraint force constants are provided in Table I.

### Curvature Analysis Using SPEX

Sampled curvature was computed using the MembraneAnalysis.jl software package (33), an implementation of the SPEX methodology (30, 31). The specific code applied was from branch ‘HPIV3_peptide’ (commit ac4d069). A target grid spacing of 15Åwas used to resolve the Fourier series coefficients. Curvature was sampled up to a *q* limit of 0.08 Å^−1^. The peptide’s position was tracked at the 23rd residue of the protein (e.g., valine 23, approximately in the middle of the sequence). An example script is available in the Zenodo repository alongside the simulation data. The quantity ⟨*J*⟩ *A*_sim_ is then reported, which is related to the spontaneous curvature *J*_0_ as

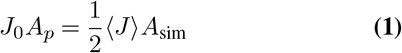

The supplemental relates ⟨*J* ⟩ *A*_sim_ to the curvature induced on a vesicle, per peptide monomer or trimer, if the spontaneous curvature is spread over a circular area (a *cap*).

### Virus Growth and Purification

Recombinant viruses were generated by reverse genetics as previously described (38, 39) using an HPIV3 clinical isolate (CI-1) (40, 41) background harboring single mutations in F (E108 or E108K) and a recombinant eGFP or mCherry cassette between genes P and M. Resulting viruses were propagated using the HAE EpiAirway AIR-100 system (MatTek Corporation). Viruses were titered by limiting dilution infection of Vero cells (ATCC), and infected cells were quantified using a Cytation5 (Agilent). All recombinant viruses were sequenced using metagenomic nextgeneration sequencing (42).

### Erythrocyte Fragment Membrane Preparation

Red blood cell fragments were used as host target membranes. Red blood cells (de-identified samples from New York Blood Center) were separated from the plasma with low-speed centrifugation (800 rcf for 5 min) and then washed at least 2 times with Dulbecco’s phosphate-buffered saline (DPBS) (Gibco). The purified red blood cells were suspended in 10% DPBS medium and stored at 4°*C* for up to 5 days. For use in cryo-electron tomography (cryo-ET) experiments, red blood cells were extruded with 10 passes through a 400 nm filter and then subsequently through a 100 nm filter using the Avanti lipid extruder. These host erythrocyte (RBC) membrane vesicles were used fresh within 24 hours of preparation.

### Cryo-electron Tomography Preparation of HPIV3 Bound to Host Target Erythrocyte Fragments

Lacey carbon gold grids, containing a continuous layer of thin carbon (Ted Pella), were plasma cleaned with a Gatan Solarus plasma cleaner for 25 seconds with an H_2_O_2_ gas mixture. 8µL drops containing 100µg*/*mL of the anti-HPIV3 HN antibody were incubated on the grids for 10 minutes and the grids were washed with DPBS to remove unadsorbed antibodies. Next, grids were blotted and placed face-down in the cell supernatant fluid, contained in a 6-well plate. Grids were blotted and placed face down in the Vero cell supernatant fluid containing CI −1− HN_H552Q*/*T193A_ in combination with host erythrocyte (RBC) membrane vesicles in a 6-well plate. Plates were incubated for 30 min at 37°C with rocking. Grids were then washed in a DPBS solution (Gibco). Grids were then plunge frozen in liquid ethane using a Vitrobot (Mark IV; Thermo Fisher Scientific Co.). For a negative control, we applied an antibody specific for measles H to the grids, and we did not observe viral particles on these grids.

### Cryo-electron Tomography Data Collection and Image Processing

Vitrified grids were imaged using a Titan Halo 300 kV transmission electron microscope (Thermo Fisher Scientific Co.), equipped with a Gatan K3 direct detector and no energy filter. Images were captured at a magnification of 18000x, giving a pixel size at the specimen level of 1.71Åfor CI-1-HN_H552Q/T193A_ and erythrocyte fragment sample. Images were acquired with SerialEM software (43) with a 3.5-5.0µm defocus and a bidirectional tilt-series of 3° steps starting from -9° to 54° and then -12° to -54°. Each tilt had a dose of 100 e −*/*Å^2^ resulting in a total dose of 2.8 e − */*Å^2^ for the tilt series. All micrograph movies were aligned using WarpEM (44). Tilt series were aligned with AreTomo (45) and then the aligned parameters were used to reconstruct the CTF corrected tomograms in WarpEM (44, 46). Final tomograms were binned by 8 giving a pixel size at the specimen level of 13.68 Åfor CI-1-HN_H552Q/T193A_ and host erythrocyte (RBC) membrane vesicles.

### HPIV3 Fusion Peptide Structure Determination Using AlphaFold

The three-dimensional structure of the HPIV3 F HRN region and FP (80 amino acids; FFGGVIGTIALGVATSAQITAAVALVEAKQARSDIEKLKE AIRDTNKAVQSVQSSIGNLIVAIKSVQDYVNKEIVPSIAR) was predicted using AlphaFold 3 (47) via the public server with default parameters.

## Results

### Surface-Parallel FP Induces Positive Curvature

The key determinant of membrane curvature generation by the hydrophobic insertion (wedging) mechanism is the position of the amphipathic helix relative to the neutral surface (17, 48). To place our results in the context of this model, we first developed a qualitative picture of how lipid structure is changed by helix insertion and depth relative to the interfacial chemistry of the proximal leaflet.

Representative molecular snapshots of the FP inserted in the proximal leaflet are shown in Fig. 2A-B, illustrating the FP positioning between the neutral surface (black) and the phosphate plane (tan). When the FP inserts deeper than the lipid headgroup region (Fig. 2A), surrounding lipids (dark blue) reorganize with both lipid headgroups and tails surrounding the FP. In contrast, when the FP is positioned at the headgroup region (Figure 2B), lipid tails concentrate beneath the FP while headgroups are displaced outward (dark blue), resulting in greater exposure of the FP to the aqueous environment. To evaluate these two lipid configurations quantitatively, we calculated the average lipid tail and headgroup orientations as a function of distance perpendicular to the FP (Fig. 2C). Positive lipid height values correspond to the proximal leaflet whereas negative values correspond to the distal leaflet. Within the proximal leaflet, lipid headgroups are displaced by FP insertion, while lipid tails conform to the shape of the FP. In contrast, lipids farther from the FP adopt an orientation aligned with the membrane normal, consistent with bulk (unperturbed) bilayer organization. The FP adopts an amphipathic insertion, with hydrophobic residues located within the bilayer core and polar residues positioned near the lipid headgroups. In the distal leaflet, lipids adjacent to the FP extend beyond the bilayer midplane (z=0), resulting in local negative curvature of the distal leaflet (Fig. 2C).

**Fig. 2.**
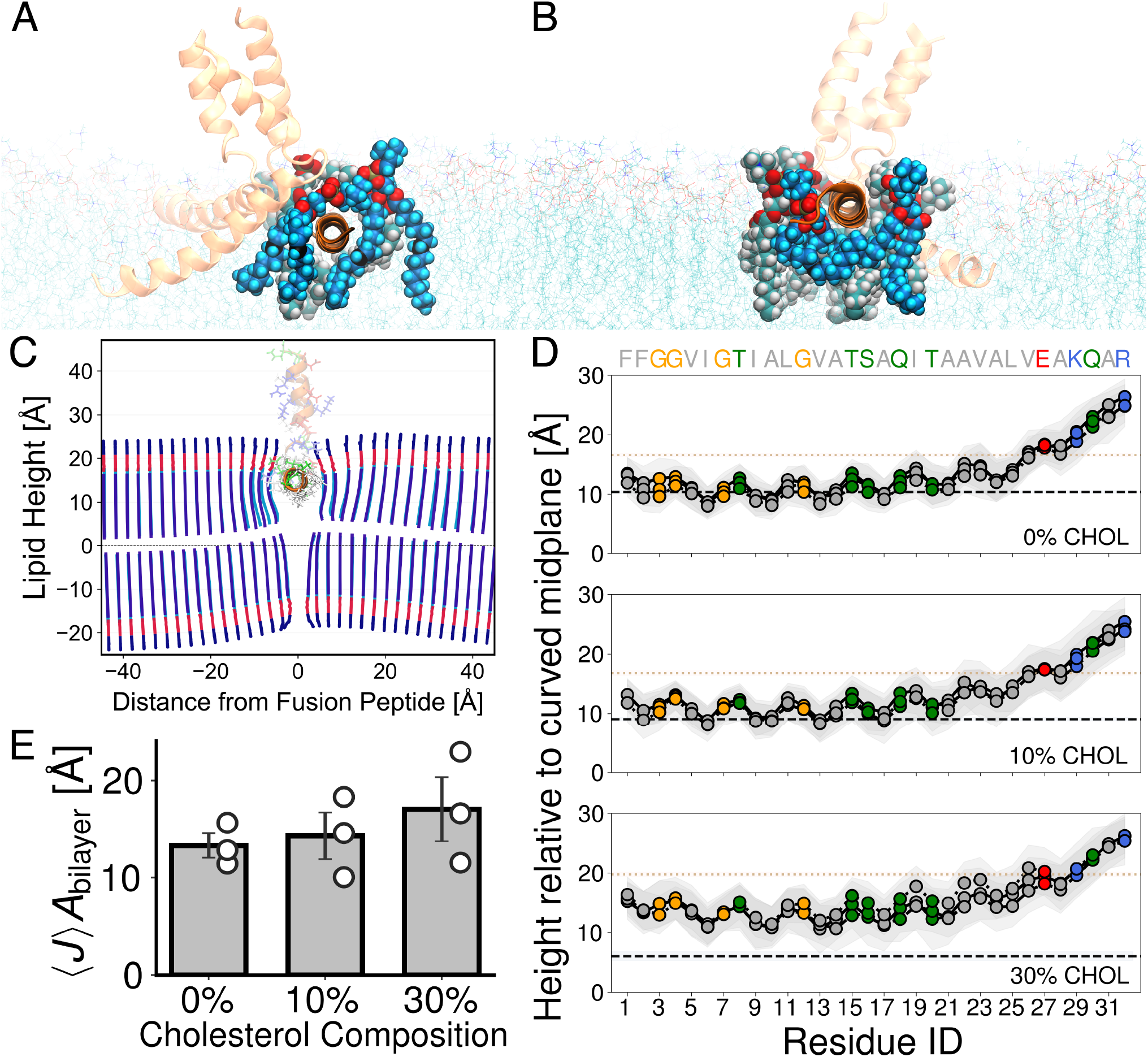
FP insertion depth, lipid orientation and membrane curvature. (A) Representative molecular snapshot of the FP in the surface-parallel configuration. The peptides (FP and adjacent N-terminal heptad repeat (HRN)) are shown in orange. The FP corresponds to the membrane-inserted segments (horizontal), while residues extending beyond this region belong to the HRN (vertical). Highlighted lipids are shown in dark blue (hydrogens cyan, phosphate tan, oxygen red, neutral surface black), with surrounding lipids in turquoise (hydrogens white, phosphate and oxygen tan and red, respectively). One FP chain inserts beneath the lipid headgroup region, with surrounding lipids reorganizing around the FP. (B) Representative molecular snapshot of a second FP chain positioned at the headgroup region, with lipid tails packing beneath the FP and proximal leaflet headgroups displaced outward.(C) Average lipid tail and headgroup orientations as a function of perpendicular distance from the FP, showing displacement of headgroups and curving of tails around the FP. Headgroups are shown in dark blue, phosphate region in red and acyl chains in blue. (D) Average FP insertion depth relative to the curved bilayer midplane (z = 0). Residues 1-26 correspond to the FP, with subsequent residues belonging to the HRN. The neutral surface (dashed line) and phosphate plane (dotted line) are shown for reference, indicating that most FP residues lie above the neutral surface across all systems (0%, 10% and 30% cholesterol). Residues are colored as follows: hydrophobic (gray), glycine (yellow), polar (green), acidic (red), and basic (blue). Gray shading corresponds to the pooled standard deviation.(E) Mean membrane curvature multiplied by the bilayer area (⟨*J*⟩*A*_bilayer_) for each system. Circles denote individual replicate means, bars indicate the mean value across replicates, and error bars represent the standard error of the mean.

We next quantified the average insertion depth of each residue in the FP relative to the curved bilayer midplane (Fig. 2D). The neutral surface, indicated by a dashed line, provides a reference for shallow (above) or deep (below) FP insertion. The neutral surface is located at 10.4, 9.1, and 6.1Åabove the bilayer midplane for 0%, 10% and 30% cholesterol, respectively. Across the FP, most residues are located above the neutral surface in all systems, indicating a predominantly shallow insertion geometry, with the greatest separation from the neutral surface observed in the 30% cholesterol system. Within the amphipathic wedging model, this insertion geometry is associated with the generation of positive membrane curvature (17, 48).

To quantify membrane curvature associated with FP insertion, we calculated the curvature generated for each system (Fig. 2E). All systems exhibit positive curvature (⟨*J* ⟩ *A*_bilayer_) with values of 13.3±1.2 Å, 14.3 ±2.4 Å, and 17.0 ±3.3Å(mean ± SEM) for 0%, 10%, and 30% cholesterol, respectively. Curvature values did not differ significantly across the three cholesterol compositions despite differences in neutral-surface position (linear regression of ⟨*J* ⟩*A*_bilayer_ vs. cholesterol concentration, slope = +0.126Åper percentage of cholesterol, t(7) = 1.19, p = 0.271, n = 9) Figs. S10 and S11.

### Transmembrane FP Induces Positive Curvature

In the proposed transmembrane insertion configuration, a key feature is lipid tail splay in FP-adjacent lipids in the distal leaflet. To evaluate tail splay, we examined lipid structural changes associated with transmembrane FP insertion (Figure 3).

**Fig. 3.**
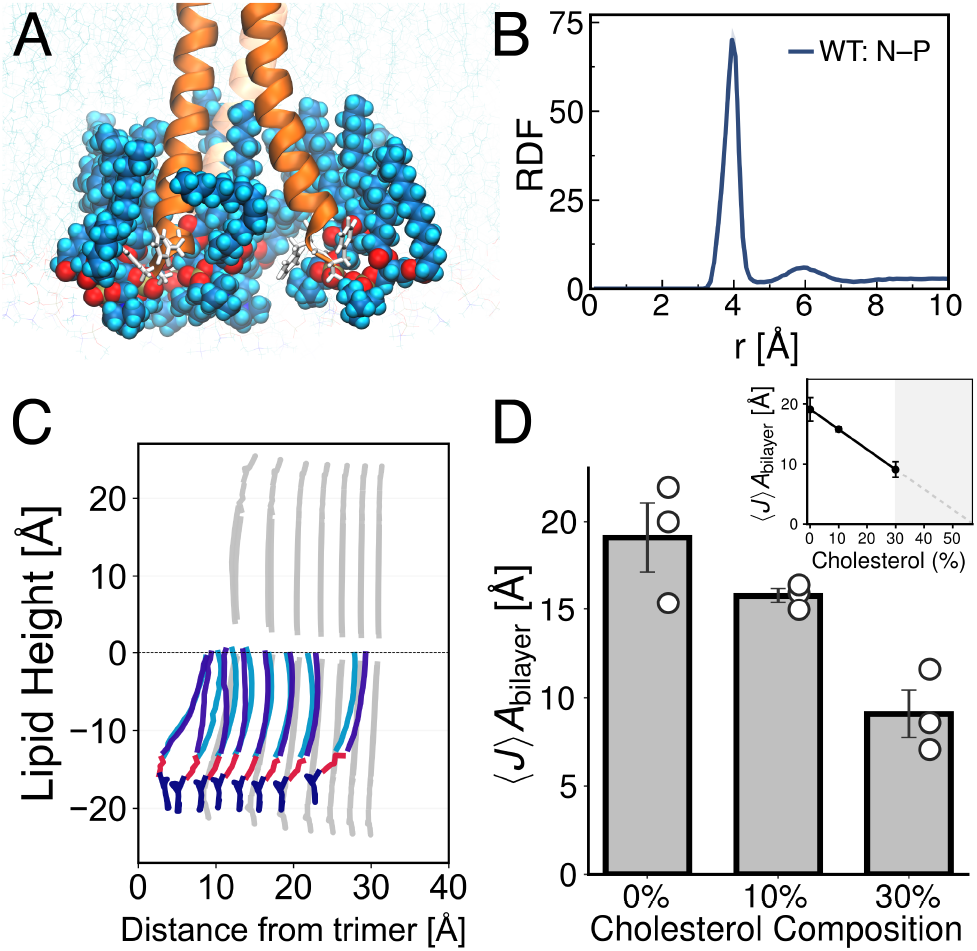
FP-induced lipid organization and membrane curvature in the transmembrane FP configuration. (A) Representative molecular snapshot of the transmembrane FPmembrane system. FPs are shown in orange. Lipids within 4.75Åof the N-terminus are shown in van der Waals representation, with surrounding lipids shown in line representation (colors as in Figure 2). FPs adopt a transmembrane orientation, with adjacent distal leaflet lipids exhibiting tail splay. (B) Radial distribution function between the N-terminal nitrogen and lipid phosphate atoms, show a strong local enrichment of lipid phosphate groups near the positively charged N-terminus. (C) Average lipid tail and headgroup positions as a function of distance from the FP (lipid profile) in the 30% cholesterol system. Gray lines show radially averaged profiles, whereas colored lines correspond to lipids within 4.75Åof the backbone N atom of the first FP residue, showing local tail splay and inward headgroup displacement in the immediate vicinity of the FP. Lipid headgroups are shown in dark blue, phosphate region in red, and acyl chains in blue. (D) Mean membrane curvature multiplied by the bilayer area ⟨*J*⟩*A*_bilayer_ for 0%, 10%, and 30% cholesterol systems. Bars indicate the mean value averaged over time and replicates, circles denote individual replicate mean, and error bars represent the standard error of the mean. The inset shows a linear regression of mean ⟨*J*⟩*A*_bilayer_ as a function of cholesterol composition, predicting zero membrane curvature at 57% cholesterol.

A representative molecular snapshot shows the FP trimer adopting a transmembrane orientation in 30% cholesterol (Figure 3A), with the N-terminus located in the distal leaflet. The surrounding distal-leaflet lipids undergo localized rearrangement around the inserted FP N-terminus, with lipid tails curving around the FP and headgroups oriented toward the N-terminus. These structural changes are concentrated near Phe1 and Phe2 (residues F110 and F111 in the native HPIV3 F sequence), which insert into the glycerol-acyl chain interface of the distal leaflet. To quantify the spatial organization of lipids around the FP N-terminus, we calculated the radial distribution function (RDF) between the N-terminal nitrogen atom and lipid phosphate groups (Fig. 3B). The distribution exhibits a pronounced first peak at approximately 3.95Åand reaches a minimum at *r* ≈ 4.75 Å, which was used to define the first lipid shell adjacent to the FP N-terminus. Lipids within this shell are directly interacting with the FP N-terminus and were used to calculate the colored lipid profiles shown in Fig. 3C. These profiles reveal the largest density of directly interacting lipids among all evaluated systems, with interactions extending farther from the trimer center than in any other FP orientation or membrane composition (Fig. 3C, Fig. S5). Individual lipid profiles also exhibit varying degrees of tail splay. To extend the analysis beyond the first lipid shell, radially averaged lipid profiles were also calculated as a function of distance from the trimer center (gray lines in Fig. 3C). Unlike the colored profiles, which are restricted to lipids directly contacting the FP N-terminus, the radially averaged profiles include both interacting and noninteracting lipids over a broader membrane region. In the radially averaged profiles (gray), the first radial shell (*x* ≈ 10 Å) shows outward displacement of lipid tails and inward displacement of headgroups toward the bilayer midplane (*y*≈ −22 Å), relative to bulk lipids (*y* ≈ −25 Å). These perturbations are confined primarily to the distal leaflet, while proximal-leaflet headgroups display only a slight upward displacement. The magnitude of the perturbation decreases with increasing distance from the FP, approaching bulk membrane organization by approximately *x* ≈ 30 Å.

To determine whether these changes in lipid organization were accompanied by changes in membrane curvature, we quantified membrane curvature for each system (Figure 3D). All three membrane compositions exhibited positive curvature (⟨ *J* ⟩*A*_bilayer_), with values of 19.1 ±1.9 Å, 15.8± 0.4 Å, and 9.1 ±1.3 Å(mean ±SEM) for membranes containing 0%, 10%, and 30% cholesterol, respectively. Curvature decreased monotonically and significantly with increasing cholesterol concentration (linear regression, slope = -0.334Åper cholesterol percent, p = 8.0 × 10^−4^, R^2^ = 0.818, n = 9; Figure 3D, inset). The linear fit intersects zero curvature at 57.2% cholesterol (95% CI: 23.0-96.2%), though this estimate requires extrapolation well beyond the tested 0-30% cholesterol range and should be interpreted accordingly (Fig. S10).

### Contribution of N-terminus Aromatics and Charge to Transmembrane FP-Induced Curvature

Having established that the FP transmembrane configuration induces positive curvature, we next examined whether specific N-terminus residues contribute to this curvature. To assess this, we generated two modified systems from the WT sequence: one in which Phe1 and Phe2 were substituted for alanine (F1A/F2A) to test the contribution of the N-terminal phenylalanines, and a second in which the N-terminus was neutralized (NNT) to isolate the contribution of N-terminal electrostatic interactions.

A representative molecular snapshot of the F1A/F2A FP is shown in Fig. 4A in 30% cholesterol. Among the lipids within the first shell surrounding the FP N-terminus (dark blue), the left lipid curves around the FP, whereas the right lipid adopts a more upright orientation. The corresponding colored lipid profiles reveal fewer directly interacting lipids and reduced tail splay than in the WT system (Fig. 4B). The radially averaged lipid profiles (gray), which include both interacting and noninteracting lipids, exhibit a smaller perturbation of distal-leaflet lipid organization than in the WT system, with a reduced displacement of lipid headgroups and tails in the first radial shell. As in the WT system, the magnitude of the perturbation decreases with increasing distance from the FP, approaching bulk bilayer organization.

**Fig. 4.**
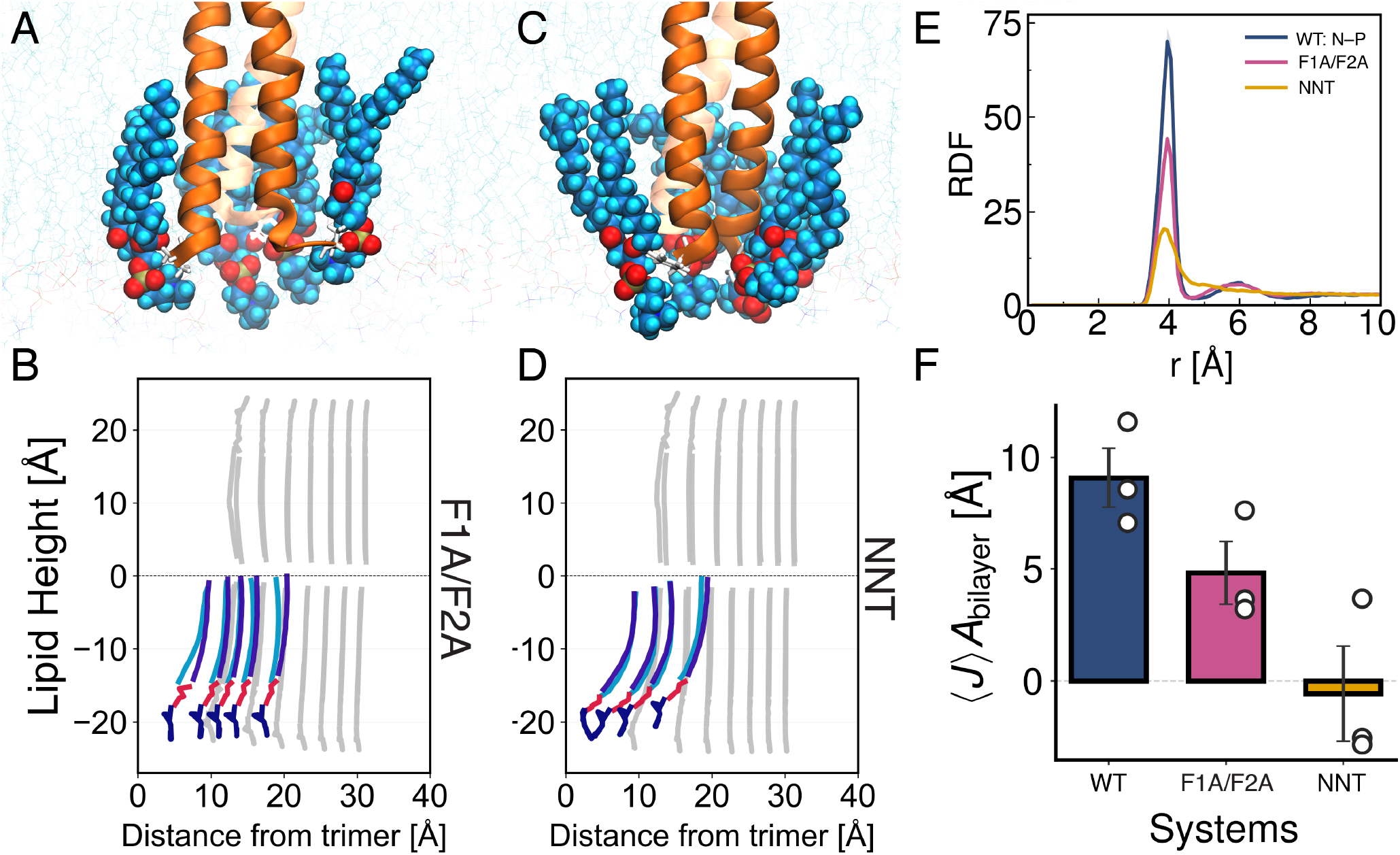
N-terminal modifications alter lipid organization and membrane curvature. (A) Representative molecular snapshot of the F1A/F2A system in 30% cholesterol. FPs are shown in orange, highlighted lipids are shown in dark blue with surrounding lipids in turquoise (colors as in Figure 2). (B) Average lipid tail and headgroup orientations as a function of distance from the FP for the F1A/F2A system, showing tail splay with headgroup positions approaching bulk values with increasing distance from the FP. Lipid headgroups (dark blue), phosphate region (red) and acyl chains (blue). (C) Snapshot of the NNT system in 30% cholesterol (lipid coloring as in panel A). (D) Average lipid tail and headgroup orientations as a function of distance from the FP for the NNT system, with minimal differences in headgroup position relative to the bilayer midplane compared with bulk lipids. (E) Radial distribution function between the N-terminal nitrogen and phosphate atoms, shows a progressive reduction in the peak amplitude from WT to F1A/F2A to the NNT system. (F) Mean membrane curvature multiplied by the bilayer area ⟨*J*⟩*A*_bilayer_ for each system, shows a progressive decrease in curvature magnitude from WT to F1A/F2A to NNT. Bars indicate the mean value averaged across replicates, circles denote individual replicate mean and error bars represent the standard error of the mean.

The NNT system exhibits a distinct lipid organization (Fig. 4C-D). In the molecular representation (Fig.4C), first-shell lipids adjacent to the FP N-terminus (dark blue) adopt a tilted configuration, with headgroups oriented toward the FP N-terminus and tails extending outward in the 30% cholesterol system. The corresponding colored lipid profiles (Fig. 4D) further illustrate that only a small population of lipids directly interacts with the FP N-terminus. These interacting lipids display a more tilted configuration than those in the F1A/F2A system, with a degree of tail splay comparable to that observed in WT. In contrast, the radially averaged lipid profiles (gray) exhibit minimal variation in lipid height with radial distance, with lipids at *x* ≈ 10Åand *x >* 30Åboth displaying heights of approximately −23 Å.

Consistent with the reduced population of directly interacting lipids observed in the colored lipid profiles, we quantified N-terminal interactions with lipid phosphate groups by calculating the radial distribution function (RDF) between the N-terminal nitrogen and phosphate atoms (Fig. 4E). All systems contain a primary peak at approximately *r* ≈ 3.9, Å, indicating close N–phosphate proximity. The peak position remains similar across systems, whereas the peak amplitude decreases progressively from WT (max *g*(*r*) = 70.12 at *r* = 3.95, Å) to F1A/F2A (*g*(*r*) = 44.28 at *r* = 3.95, Å) to NNT (*g*(*r*) = 20.45 at *r* = 3.85, Å). This corresponds to an approximately threefold reduction in local phosphate density between WT and the NNT system. A similar progressive decrease in peak amplitude was observed in the 0% cholesterol simulations (Fig. S4). To evaluate differences in membrane curvature across the WT, F1A/F2A, and NNT FP systems, we quantified curvature (⟨*J*⟩ *A*_bilayer_) for each modified system in 30% cholesterol (Figure 4F). Similar to the WT system, the F1A/F2A system also induced positive curvature (Figure 4F), whereas the NNT system exhibited near-zero curvature with a negative mean. The magnitude of curvature decreased across the three systems, with values of 9.1 ±1.3 Å, 4.8 ±1.4 Å, and − 0.5 ±2.1Å(mean ±SD) for the WT, F1A/F2A, and NNT systems, respectively. However, these differences did not reach statistical significance at 30% cholesterol (F(2,15) = 2.34, p = 0.138). In cholesterol-free membranes, the effect of the N-terminal modifications differed, with a significant difference among the three systems (F(2,8) = 13.60, p = 0.0027; Fig. S8). F1A/F2A generated significantly lower curvature than both WT and NNT (p = 0.0023 and p = 0.0210, respectively), whereas WT and NNT did not differ significantly (p = 0.278). The NNT system exhibited greater curvature than the F1A/F2A system despite the same ordering of N-phosphate interactions (Fig. S2).

### Membrane Curvature Depends on HPIV3 FP Oligomeric State and Configuration

To compare membrane curvature induced by the HPIV3 FP monomer and trimer, we quantified curvature generated for both systems under identical membrane compositions and orientations (Fig. S1). Across all conditions evaluated, trimers generated greater curvature than monomers, but the increase was not proportional to the number of peptides. For transmembrane systems, mean curvature was 8.0 ±0.9Åfor the monomer and 15.8 ±0.4Åfor the trimer at 10% cholesterol, and 1.2 ± 3.7Åand 9.1 ± 1.3 Å, respectively, at 30% cholesterol (mean ± SEM). A comparable difference was observed for the surface-parallel orientation, with monomer and trimer curvature of 7.4 ± 3.6Åand 14.3 ± 2.4Åat 10% cholesterol and 13.2 ± 5.2Åand 17.0 ± 3.3Åat 30% cholesterol. Beyond the monomer-trimer comparison, the two-way ANOVA identified significant main effects of both cholesterol concentration and system on membrane curvature (Figs. S6 and S7).

We next compared membrane curvature generated by WT trimers in the transmembrane and surface-parallel orientations across cholesterol compositions (Fig. S2). At 0% cholesterol, the transmembrane FP orientation generated greater curvature than the surface-parallel FP orientation, with mean values of 19.1 ±2.0 and 13.3 ±1.2 Å, respectively. At 10% cholesterol, curvature was similar between the two orientations, with values of 15.8 ±0.4 and 14.3± 2.4Åfor the transmembrane and surface-parallel systems, respectively. At 30% cholesterol, however, the surface-parallel orientation generated greater curvature than the transmembrane orientation, with values of 17.0 ±3.3 and 9.1± 1.3 Å, respectively. None of these three pairwise differences reached statistical significance individually (Welch’s t-test; 0%: p = 0.078; 10%: p = 0.603; 30%: p = 0.124), consistent with the modest sample size of (n = 3) replicates per condition. However, this apparent crossover is consistent with the significant System × Cholesterol interaction established by two-way ANOVA across all six systems (F(6,28) = 4.31, p = 0.0034, Table IV). Curvature declined for the transmembrane orientation while increasing for the surface-parallel orientation. These opposing trends are likely contributors to the interaction and indicate that cholesterol’s effect on curvature depends on FP orientation rather than acting uniformly.

Peptide Number and Geometry were additionally evaluated separately at 0% and 30% cholesterol (Fig. S8). At 0% cholesterol, Peptide Number could not be evaluated because only trimer systems were simulated, and the effect of Geometry was not significant (p = 0.262). At 30% cholesterol, where all four Peptide Number × Geometry combinations were simulated, Geometry was significant (p = 0.0009), whereas Peptide Number was not significant (p = 0.886). In the pooled analysis, both Peptide Number and Geometry were significant (p = 0.005 and p = 0.002, respectively). However, after stratification by cholesterol, Peptide Number was not significant at 30% while Geometry remained significant at 30% (Fig. S9).

### Cryo-Electron Tomography Reveals Membrane Curvature at Sites of HPIV3-Host Cell Contact

Cryo-ET was performed on HPIV3 particles bound to host cells to characterize membrane curvature at sites of virus-cell contact. Representative tomograms show localized deformation of the host-cell membrane adjacent to bound viral particles (Fig. 5A). Host-cell membrane curvature was quantified within a ±20nm region at the virus-cell interface. Local curvature values were mapped onto the membrane contour, revealing localized regions of curvature in the host-cell membrane immediately adjacent to bound viral particles (Fig. 5A). Across all analyzed particles from the clinical isolate, the curvature distribution was shifted toward positive values, with a mean curvature of *µ*= 1.80 ×10^−4^Å^−1^ (95% CI : [1.3927, 2.2325] × 10^−4^Å^−1^)(Fig. 5B, top). Overall, 63.9% of measured membrane regions exhibited positive curvature, whereas 36.1% exhibited negative curvature. In contrast, the HN-P111S variant, used as a negative control in which FP insertion is absent, exhibited a curvature distribution shifted toward negative values, with a mean curvature of *µ*= −0.96 ×10^−4^Å^−1^ (95% CI : [−2.0405, −0.2605] ×10^−4^Å^−1^) (Fig. 5B, bottom). Overall, 55.4% of measured regions exhibited negative curvature, compared with 44.6% exhibiting positive curvature. Bootstrap analysis further supported a difference in mean curvature between the two conditions. The mean curvature of the clinical isolate was higher than that of the HN-P111S variant by 2.76 × 10^−4^ Å^−1^ (bias corrected accelerated (BCa) 95% CI: [1.95,, 3.89] × 10^−4^ Å^−1^ ; Table V). In contrast, the BCa 95% confidence interval for the difference in variance included zero, indicating no supported difference in variance between the two groups. Together, these measurements show that positive host-cell membrane curvature is enriched at sites of HPIV3 attachment under conditions that permit FP insertion, whereas the non-inserting control is shifted toward negative curvature.

**Fig. 5.**
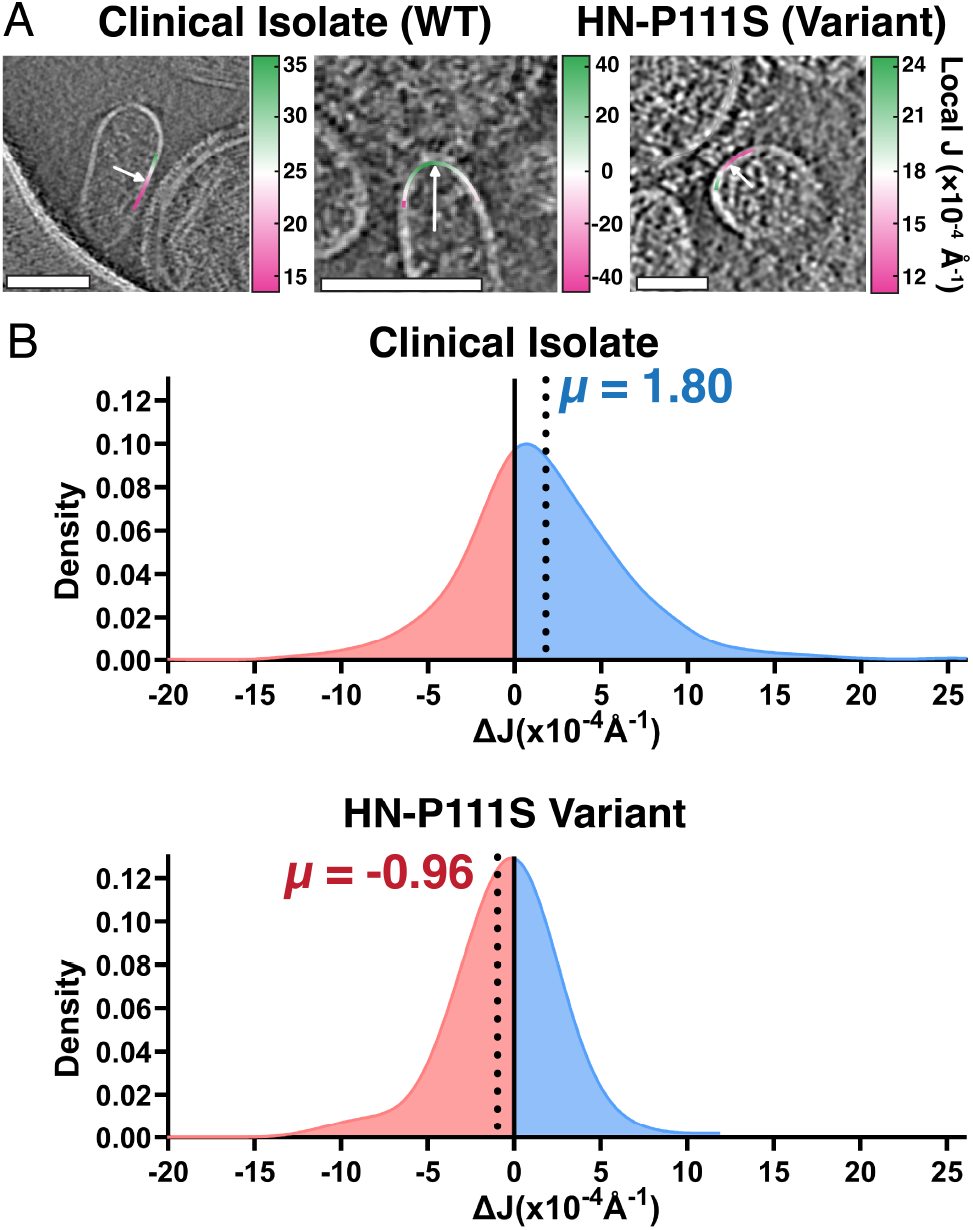
Curvature analysis of host-cell membranes at HPIV3–host contact sites. (A) Representative cryo-electron tomography slices showing HPIV3 particles interacting with host red blood cell membrane fragments for the clinical isolate (left and right) and HN-P111S variant (center). Local membrane curvature along the traced host-cell membrane is indicated by the green-to-magenta colormap, with green representing lower curvature and magenta representing higher curvature. White arrows indicate the point of closest contact between the host-cell and viral membranes. The corresponding Δ*J* value is shown below each image. (B) Kernel density estimates of Δ*J* for all analyzed host-cell membrane regions for the clinical isolate (top) and HN-P111S variant (bottom). Dashed blue lines indicate the mean Δ*J*. Red and blue shading indicate negative and positive Δ*J* values, respectively, with the percentage of measurements in each region indicated. Scale bars, 100 nm.

## Discussion

Molecular simulations demonstrate that both HPIV3 FP surface-parallel and transmembrane configurations induce positive curvature. In the surface-parallel configuration, shallow FP insertion near or above the proximal-leaflet neutral surface generates positive curvature through the hydrophobic insertion (wedging) mechanism (17). In contrast, the transmembrane FP configuration deforms the *distal* leaflet, generating local distal-leaflet negative curvature. Because curvature is defined here from the perspective of the fusion protein, which extends above the proximal leaflet, this deformation is reported as positive curvature. Through *in silico* mutations and structural information (lipid traces and radial distribution functions), lipid interactions with the bulky and positively charged N-terminus are identified as the molecular mechanism for transmembrane FP-induced curvature. As discussed in detail below, cholesterol content and the number of FPs (monomer or trimer) influence the magnitude of curvature. These simulation results are consistent with membrane curvature observed experimentally by cryo-ET. Relative to the mutant system in which the F protein is not inserted into the host membrane, cryo-ET reveals a shift toward positive host-membrane curvature at HPIV3-cell contact sites, consistent with the positive curvature generated by both simulated configurations.

### Surface-parallel FP induces positive curvature through wedging

The hydrophobic wedging mechanism (17, 48–50) predicts that amphipathic helix insertion depth determines the sign and magnitude of curvature. Shallow FP insertion above the proximal-leaflet neutral surface generates positive curvature, whereas below the neutral surface curvature generation is negative (17, 48). Consistent with this model, most HPIV3 FP residues occupy the proximal-leaflet interfacial region near or above the neutral surface across all cholesterol compositions (Fig. 2D). Above the neutral surface, lipid headgroups are displaced outward while their tails pack beneath the FP (Fig. 2B). Below the neutral surface, headgroups would instead move above the FP and tails would be pushed out — a structural motif we do not observe here. The FP’s position relative to the neutral surface varies with cholesterol content, lying progressively farther above it as cholesterol increases. This greater separation reflects both a modest reduction in FP insertion depth and a shift of the neutral surface toward the bilayer midplane (Fig. 2D), changes consistent with the known effects of cholesterol on lipid packing and ordering (51). Positive curvature magnitude increased modestly as the FP moved higher above the neutral surface with cholesterol, although the differences among compositions were not statistically significant (Fig. 2E). Together, the shallow position of the FP relative to the neutral surface and the observed positive curvature provide convergent support for a hydrophobic insertion (wedging) mechanism in the surface-parallel configuration.

### Transmembrane FP generates curvature through distal-leaflet reorganization

Positive curvature generation by the transmembrane FP configuration in the all-atom model is reported (Fig. 3D) alongside lipid reorganization in the distal leaflet around the FP N-terminus (Fig. 3C). This reorganization is concentrated near Phe1 and Phe2 (residues F110 and F111 in the native HPIV3 F sequence), which reside at the glycerolacyl-chain interface of the distal leaflet (Fig. 3A,C). Lipid traces indicate those lipids interacting with the N-terminal amine display characteristics consistent with negative distal leaflet curvature, with headgroups moving in toward the FP (Fig. 3C). This inward headgroup movement is accompanied by a pronounced short-range peak in the N-terminal nitrogen-phosphate RDF, reflecting local enrichment of lipid phosphate groups around the N-terminus (Fig. 3B). A similar effect was reported for the influenza fusion peptide, in which partial burial of the charged N-terminal amino group in the membrane-spanning configuration promoted local lipid tilting (52). Membrane cholesterol further modulates this behavior (Fig. S5), strongly reducing the magnitude of curvature generated by the transmembrane FP configuration (Fig. 3D). These observations support a mechanism in which local lipid reorganization in the distal leaflet contributes to negative curvature, which (when retaining the proximal leaflet definition of the sign of curvature) is reported as positive curvature. Weakening of the effect by cholesterol suggests that local lipid ordering counteracts the deformation of the transmembrane configuration. This effect appears systemand configurationdependent: for HIV GP41, increased lipid order was proposed to enhance curvature (53), whereas for the HPIV3 transmembrane FP increasing cholesterol content reduces curvature, while the surface-parallel configuration shows a slight increase with cholesterol that is not statistically significant.

### Phe1/Phe2 and N-terminal charge contribute to transmembrane curvature generation

The F1A/F2A system was designed as a test of the contribution of Phe1/Phe2 side-chain bulk to lipid reorganization and distal-leaflet curvature generation, while preserving hydrophobicity at these positions. In the 30% cholesterol membrane, replacing Phe1 and Phe2 with alanine reduced local phosphate enrichment around the N-terminus and lowered mean curvature relative to WT (Fig. 4B,E–F). The weaker perturbation in the radially averaged F1A/F2A lipid profiles further suggests that Phe1 and Phe2 contribute to the collective reorganization of distal-leaflet lipids around the inserted N-terminus (Fig. 4B). Mutational analysis of the PIV5 in Ref. (54) previously identified the N-terminal phenylalanine F103 as important for membrane perturbation.

The NNT system was designed as a test of contribution of the electrostatic interactions between the positively charged N-terminus and lipid phosphate groups to distal-leaflet curvature generation. In the 30% cholesterol membrane, neutralizing the N-terminus produced the largest reduction in local phosphate enrichment and reduced mean curvature to approximately zero (Fig. 4E–F). The small population of lipids directly contacting the neutral N-terminus still exhibited tail splay, whereas the radially averaged profiles showed little deviation from bulk lipid organization (Fig. 4D). The reduction in curvature with the neutral N-terminus and phenylalanine substitutions (Fig. 4F), combined with the trace of the deformed lipids (Figs. 3C, 4B, 4D) interacting with the Nterminus (Fig. 4E) is strongly suggestive of a mechanism in which lipids interacting with the N-terminus are splayed by the bulky phenylalanines, creating a lipid annulus inducing negative curvature. There are fewer N-terminal contacts *without* the phenylalanines suggesting that the smoother surface of the alanine-substituted FP favors an unsplayed lipid configuration that is less likely to interact past the terminus, whereas intuitively the smaller alanine might have allowed closer contacts with the terminus. The drop in curvature induction with cholesterol across the transmembrane configurations may be a result of a decreased sensitivity of the lipid matrix to the N-terminus-interacting annular lipids. See Figs. S10 and S11 for a consolidated presentation of the drop in curvature with cholesterol, including an illustration of the difficulty of estimating the cholesterol fraction at which curvature induction is negated by cholesterol.

### The HPIV3 FP trimer increases membrane-curvature magnitude

For both FP configurations and cholesterol concentrations (10% and 30%), trimers induce greater curvature than the monomer (Fig. S1) but less than a factor of three at the short lengthscales modeled here. Although the present simulations do not examine FP assembly, the greater curvature generated by the preassembled HPIV3 trimer complements prior PIV5 studies linking larger FP assemblies to enhanced membrane perturbation (22, 24), and is consistent with the trimeric architecture enhancing FP-driven membrane curvature (55).

### Cryo-ET supports positive host-membrane curvature at virus–host contact sites

Cryo-ET provides an experimental test of whether the positive membrane curvature observed in the simulated FP systems is also present at virus-host contact sites. Unlike the MD simulations, which isolate the effect of the FP, cryo-ET captures host-membrane deformation at intact virus-host contact sites, where the complete viral fusion machinery is present. Cryo-ET analysis of the change in curvature (Δ*J*) at virus-cell contact sites the clinical isolate revealed a distribution shifted toward positive values (Fig. 5), consistent with simulation predictions of both surface-parallel and transmembrane configurations. To test whether this shift was associated with the inserted FP, the clinical isolate was compared with the non-inserting HN-P111S variant. Mean Δ*J* was significantly higher in the clinical isolate than in the HN-P111S variant, whereas no difference in variance was supported (Table V). The change from predominantly positive to predominantly negative Δ*J* in HN-P111S is consistent with the membrane-inserted FP contributing to positive host-membrane curvature, rather than simply increasing curvature variability.

To relate experimental Δ*J* to curvature calculated in the simulations, the Supplemental Material derives a simple estimate of curvature induction on a vesicle by a circular region (a *cap*) with spontaneous curvature *J*_0_ as derived from input *J* ⟨*A* ⟩. Figures 2-4 report the values of *J* ⟨*A* ⟩ that can be translated into such a cap spontaneous curvature, equivalent to the effect of the fusion peptide near the virus-host adhesion site. The cryo-ET derived shift in the mean, approximately 2 × 10^−4^ Å^−1^, is considerably larger than the curvature induced by the trimer of FPs of a fusion protein according to our modeling estimate, using (Eq. S38):

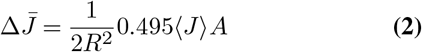

where *J* ⟨*A*⟩ is approximately 20 Å. For a vesicle of radius 50 nm, simulations predict curvature induction to be 2 × 10^−5^ Å^−1^. This difference may arise because multiple FPs are responsible for inducing curvature over the observation region in the cryo-ET data, the single spherical harmonic employed resulting in Eq. 2 may not be consistent with the image curvature extraction, the simulation may underestimate curvature induction, or additional curvature-inducing mechanisms may be involved. Figure S3 of the Supplemental Material shows that the curvature coupling lengthscale accessible to simulation may indeed underestimate values at very long lengthscales. Extrapolating the length-scale-dependent curvature to the macroscopic scale increases its magnitude by approximately a factor of two. Considering the unknown number of FPs inducing curvature within the 20 nm window and the gap in lengthscales between simulation and experiment, the qualitative agreement of cryo-ET and simulation points to the FP component as substantially changing the shape of the host membrane, supporting the formation of a bump (see Fig. 6 and discussion below).

**Fig. 6.**
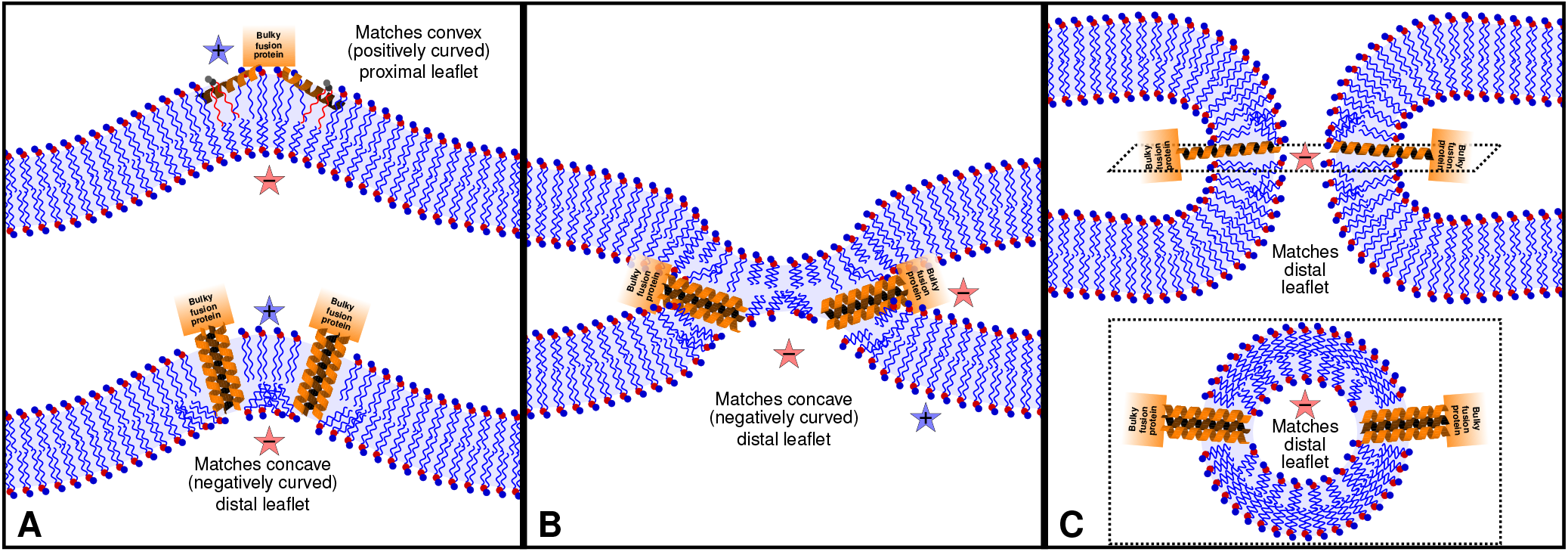
Proposed model for the role of distinct HPIV3 FP insertion states during HPIV3 viral membrane fusion. The host membrane transitions from the initial contact state, protrusion (12) (panel A) to a hemifusion diaphragm (B), and finally to fusion pore formation (C) (12, 13). In panel A, the surface-parallel (top) and transmembrane (bottom) FP configurations generate distinct membrane deformations. The surface-parallel configuration induces positive curvature (blue stars) through the hydrophobic insertion (wedging) mechanism, whereas the transmembrane configuration generates negative curvature (red stars) in the distal leaflet through local lipid deformation coupled to N-terminal interactions with lipid phosphate groups. In panels B and C are shown the proposed role of the transmembrane configuration in membrane geometries associated with hemifusion (B) and fusion pore formation (C) with positive and negative curvature labeled with stars. The hypothesized sequence of FP insertion states during membrane fusion is based on the membrane-deformation mechanisms identified in this study.

Comparison of the clinical isolate with the HN-P111S variant provides further insight into the origin of the membrane curvature observed by cryo-ET. At contact sites formed by the clinical isolate, the host-cell membrane curvature distribution was shifted toward positive values, whereas the HN-P111S variant, in which F activation and FP insertion are absent, exhibited a distribution shifted toward negative curvature. The curvature distribution for the HN-P111S variant was also narrower than that of the clinical isolate, consistent with a more uniform membrane curvature in the absence of F activation and FP insertion. In contrast, the broader distribution observed for the clinical isolate is consistent with spatially heterogeneous membrane deformation at virus–cell contact sites. This contrast supports the conclusion that the positive curvature observed at the virus–cell interface is driven by the FP, rather than arising from the native curvature of the host-cell membrane. Together, the FP-focused molecular dynamics simulations and cryo-ET measurements provide convergent computational and experimental evidence that the HPIV3 FP drives positive host-membrane curvature at intact HPIV3–cell contact sites.

### Proposed roles of HPIV3 FP insertion states during membrane fusion

The leaflet-specific curvature-inducing effect of the fusion peptide can be connected to how it would support separate stages of the fusion pathway by considering studies of the effects of lipids with known spontaneous curvature. The initial stage of fusion (contact) requires positive curvature near the contact site (Fig. 6A) and likely disruption of the merging leaflet surfaces (56). At this stage is where the comparison with cryo-ET is relevant. Following initial (unmerged) contact, stalk formation is favored by negative spontaneous curvature lipids due to the initial saddle-like structure of the stalk (57), with simulations indicating transmembrane domains of fusion proteins may favor such a stalk (58). In critical later stages of membrane fusion, including hemifusion and small fusion pores, the most highly curved regions exhibit negative curvature in *both* the proximal and distal leaflets. Specifically, the triple leaflet junction of hemifusion is concave on all three leaflets (panel B, Fig. 6), whereas, due to a quirk of saddle (negative Gaussian) curvature, *both leaflets* of the fusion pore (panel C, Fig. 6) have negative curvature when measured at the leaflet surface (see e.g. (59)). Once a small pore has formed, negative spontaneous curvature in the contacting or distal monolayer can promote pore enlargement. Thus, lipids with either positive or negative spontaneous curvature can promote or inhibit fusion depending on when and into which leaflet they are introduced (60–62). Positively curved lipids can promote pore formation in the hemifusion diaphragm when present in the distal leaflets, while inhibiting expansion of the hemifused membranes when present in the proximal leaflets. Negatively curved lipids generally have the opposite effect, favoring stalk formation and pore expansion in the proximal leaflets.

The surface-parallel and transmembrane HPIV3 FP configurations generate curvature through mechanisms centered in different leaflets, suggesting that insertion configuration is likely to influence different stages of the viral fusion pathway (Fig. 6). Here we propose that, while either configuration can support the initial (contact) phase of fusion, the transmembrane configuration can support later stages of fusion (hemifusion and full fusion). The distal-leaflet deformation associated with the transmembrane HPIV3 FP could stabilize a hemifusion-like intermediate before pore opening and contribute to negatively curved regions of the expanding pore neck after pore nucleation (63), while additional protein-generated membrane stresses or F-protein conformational changes drive progression between these FP configurations (64). These findings support a model in which HPIV3 FP insertion configuration determines how curvature is distributed across the bilayer, providing a mechanism by which the FP can influence the stability and progression of distinct membrane-fusion intermediates.

## Conclusions

This work provides a mechanistic description HPIV3 FPgenerated membrane curvature for two distinct FP insertion configurations within the host membrane. Membrane curvature induction depends strongly on FP orientation and arises through distinct membrane-deformation mechanisms. The surface-parallel conformation induces curvature through localized deformation of the proximal leaflet, consistent with the hydrophobic insertion (wedging) mechanism. The transmembrane configuration induces curvature by deforming the *distal* leaflet. FP modification analysis further revealed that the N-terminal phenylalanines and positively charged N-terminus make complementary contributions to curvature generation, demonstrating that membrane deformation depends not only on FP orientation but also on the molecular details of FP-lipid interactions. The distinct leaflets associated with the surface-parallel and transmembrane configurations further suggest that these insertion states influence different stages of the fusion process. The surface-parallel configuration generates proximal-leaflet curvature consistent with membrane protrusion models proposed to facilitate early membrane contact, whereas the transmembrane configuration generates distal-leaflet curvature compatible with membrane structures associated with hemifusion and fusion pore formation. A transition of the HPIV3 FP from surface-parallel to transmembrane could allow the FP to generate the distinct membrane deformations required at different stages of the fusion pathway. This study establishes a well-defined approach for linking molecular mechanisms to viral and host membrane shapes during fusion, using SPEX’s linear-response modeling of curvature sensitivity for quantitative comparison with static curvature observed in real viral fusion. The agreement between simulated and experimentally observed membrane curvature using cryo-ET supports the proposed role of the HPIV3 FP as a key membrane-deforming element of the viral fusion machinery, bridging atomistic descriptions of FP-induced membrane curvature with membrane deformation observed during intact virus-host interactions.

## Notes

The authors declare no competing financial interest.

## Data and Code Availability

Molecular dynamics simulation trajectories (DCD format) and the corresponding topology files generated in this study are available on Zenodo (10.5281/zenodo.22685726 (monomers), 10.5281/zenodo.22707224 (trimers), 10.5281/zenodo.22707242 (mutants)). Pre-aligned tilts for all cryoET data have been deposited on Zenodo (10.5281/zenodo.22283142). Membrane curvature analysis source code is available at https://github.com/tmarc292/CurvaTrace.

## Acknowledgments

M.B., L.J.S.L., P.S.B., A.J.S., and J.Z. and were supported by the intramural research program of the *Eunice Kennedy Shriver* National Institute of Child Health and Human Development (NICHD) of the National Institutes of Health (NIH). T.C.M., N.M., and A.M. were supported by funding from the National Institutes of Health (RO1AI175362 and U19AI181984 to A.M.) and the Sharon Golub Fund at Columbia University Vagelos College of Physicians and Surgeons. Computational resources were provided by the Biowulf high performance computing facility of the NIH, as well as by dedicated resources provided by the NICHD. This work used NAMD for molecular dynamics simulations. NAMD was developed by the Theoretical and Computational Biophysics Group in the Beckman Institute for Advanced Science and Technology at the University of Illinois at Urbana-Champaign. Cryo-ET imaging was performed at the Imaging Facility at CUNY Advanced Science Research Center. During preparation of this work, the authors used ChatGPT and Claude to assist with manuscript editing, code development, and theoretical derivations. The authors reviewed and edited all AI-assisted content and take full responsibility for the content of the publication.

## Supporting Information

The Supporting Information is available free of charge at `link`.

Detailed descriptions of the molecular dynamics and cryoelectron tomography data analysis methods, including lipid trace and radial distribution function analysis, membrane curvature analysis and statistical methods; a summary table of simulation systems and restraint conditions (Table S1); ANOVA and post-hoc comparison tables (Tables S2–S4); cryo-ET curvature statistics (Table S5); supplemental figures (Figures S1–S11); and a theoretical derivation of expected curvature deformation over a region of a spherical vesicle surface.

## References

1. Xin Wang, You Li, Maria Deloria-Knoll, Shabir A. Madhi, Cheryl Cohen, Vina Lea Arguelles, Sudha Basnet, Quique Bassat, W. Abdullah Brooks, Marcela Echavarria, Rodrigo A. Fasce, Angela Gentile, Doli Goswami, Nusrat Homaira, Stephen R. C. Howie, Karen L. Kotloff, Najwa Khuri-Bulos, Anand Krishnan, Marilla G. Lucero, Socorro Lupisan, Maria Mathisen, Kenneth A. McLean, Ainara Mira-Iglesias, Cinta Moraleda, Michiko Okamoto, Histoshi Oshitani, Katherine L. O’Brien, Betty E. Owor, Zeba A. Rasmussen, Barbara A. Rath, Vahid Salimi, Pongpun Sawatwong, J. Anthony G. Scott, Eric A. F. Simões, Viviana Sotomayor, Donald M. Thea, Florette K. Treurnicht, Lay-Myint Yoshida, Heather J. Zar, Harry Campbell, Harish Nair, and Respiratory Virus Global Epidemiology Network. Global burden of acute lower respiratory infection associated with human parainfluenza virus in children younger than 5 years for 2018: a systematic review and meta-analysis. The Lancet. Global Health, 9(8):e1077–e1087, August 2021. ISSN 2214-109X. doi: 10.1016/S2214-109X(21)00218-7.

2. Auke E. S. de Zwart, Annelies Riezebos-Brilman, Jan-Willem C. Alffenaar, Edwin R. van den Heuvel, Christiaan Tji Gan, Wim van der Bij, Huib A. M. Kerstjens, and Erik A. M. Verschuuren. Evaluation of 10 years of parainfluenza virus, human metapneumovirus, and respiratory syncytial virus infections in lung transplant recipients. American Journal of Transplantation: Official Journal of the American Society of Transplantation and the American Society of Transplant Surgeons, 20(12):3529–3537, December 2020. ISSN 1600-6143. doi: 10.1111/ajt.16073.

3. Dimpy P. Shah, Pankil K. Shah, Jacques M. Azzi, and Roy F. Chemaly. Parainfluenza virus infections in hematopoietic cell transplant recipients and hematologic malignancy patients: A systematic review. Cancer Letters, 370(2):358–364, January 2016. ISSN 1872-7980. doi: 10.1016/j.canlet.2015.11.014.

4. Matteo Porotto, Samantha G. Palmer, Laura M. Palermo, and Anne Moscona. Mechanism of fusion triggering by human parainfluenza virus type III: communication between viral glycoproteins during entry. The Journal of Biological Chemistry, 287(1):778–793, January 2012. ISSN 1083-351X. doi: 10.1074/jbc.M111.298059.

5. Rui Xu, Samantha G. Palmer, Matteo Porotto, Laura M. Palermo, Stefan Niewiesk, Ian A. Wilson, and Anne Moscona. Interaction between the hemagglutinin-neuraminidase and fusion glycoproteins of human parainfluenza virus type III regulates viral growth in vivo. mBio, 4(5):e00803–00813, October 2013. ISSN 2150-7511. doi: 10.1128/mBio.00803-13.

6. Tara C. Marcink, Gillian Zipursky, Wenjing Cheng, Kyle Stearns, Shari Stenglein, Kate Golub, Frances Cohen, Francesca Bovier, Daniel Pfalmer, Alexander L. Greninger, Matteo Porotto, Amedee des Georges, and Anne Moscona. Subnanometer structure of an enveloped virus fusion complex on viral surface reveals new entry mechanisms. Science Advances, 9(6):eade2727, February 2023. ISSN 2375-2548. doi: 10.1126/sciadv.ade2727.

7. Matteo Porotto, Zuhair W. Salah, Long Gui, Ilaria DeVito, Eric M. Jurgens, Hong Lu, Christine C. Yokoyama, Laura M. Palermo, Kelly K. Lee, and Anne Moscona. Regulation of paramyxovirus fusion activation: the hemagglutinin-neuraminidase protein stabilizes the fusion protein in a pretriggered state. Journal of Virology, 86(23):12838–12848, December 2012. ISSN 1098-5514. doi: 10.1128/JVI.01965-12.

8. Matteo Porotto, Matthew Murrell, Olga Greengard, and Anne Moscona. Triggering of human parainfluenza virus 3 fusion protein (F) by the hemagglutinin-neuraminidase (HN) protein: an HN mutation diminishes the rate of F activation and fusion. Journal of Virology, 77(6): 3647–3654, March 2003. ISSN 0022-538X. doi: 10.1128/jvi.77.6.3647-3654.2003.

9. Sally M. Kephart, Nancy Hom, and Kelly K. Lee. Visualizing intermediate stages of viral membrane fusion by cryo-electron tomography. Trends in Biochemical Sciences, 49(10):916–931, October 2024. ISSN 0968-0004, 1362-4326. doi: 10.1016/j.tibs.2024.06.012.

10. Judith M. White, Amanda E. Ward, Laura Odongo, and Lukas K. Tamm. Viral Membrane Fusion: A Dance Between Proteins and Lipids. Annual review of virology, 10(1):139–161, September 2023. ISSN 2327-056X. doi: 10.1146/annurev-virology-111821-093413.

11. Gonen Golani and Ulrich S. Schwarz. High curvature promotes fusion of lipid membranes: Predictions from continuum elastic theory. Biophysical Journal, 122(10):1868–1882, May 2023. ISSN 0006-3495. doi: 10.1016/j.bpj.2023.04.018.

12. Leonid V. Chernomordik and Michael M. Kozlov. Protein-Lipid Interplay in Fusion and Fission of Biological Membranes. Annual Review of Biochemistry, 72(1):175–207, June 2003. ISSN 0066-4154, 1545-4509. doi: 10.1146/annurev.biochem.72.121801.161504.

13. Alexandr Chanturiya, Leonid V. Chernomordik, and Joshua Zimmerberg. Flickering fusion pores comparable with initial exocytotic pores occur in protein-free phospholipid bilayers. Proceedings of the National Academy of Sciences of the United States of America, 94(26): 14423–14428, December 1997. ISSN 0027-8424. doi: 10.1073/pnas.94.26.14423.

14. Andres Chang and Rebecca E. Dutch. Paramyxovirus Fusion and Entry: Multiple Paths to a Common End. Viruses, 4(4):613–636, April 2012. ISSN 1999-4915. doi: 10.3390/v4040613.

15. Richard M. Epand. Fusion peptides and the mechanism of viral fusion. Biochimica et Biophysica Acta (BBA) - Biomembranes, 1614(1):116–121, July 2003. ISSN 0005-2736. doi: 10.1016/S0005-2736(03)00169-X.

16. Sascha Martens and Harvey T. McMahon. Mechanisms of membrane fusion: disparate players and common principles. Nature Reviews Molecular Cell Biology, 9(7):543–556, July 2008. ISSN 1471-0080. doi: 10.1038/nrm2417.

17. Felix Campelo, Harvey T. McMahon, and Michael M. Kozlov. The Hydrophobic Insertion Mechanism of Membrane Curvature Generation by Proteins. Biophysical Journal, 95(5): 2325–2339, September 2008. ISSN 0006-3495. doi: 10.1529/biophysj.108.133173.

18. Marc Fuhrmans and Siewert J. Marrink. Molecular View of the Role of Fusion Peptides in Promoting Positive Membrane Curvature. Journal of the American Chemical Society, 134(3): 1543–1552, January 2012. ISSN 0002-7863, 1520-5126. doi: 10.1021/ja207290b.

19. Beatriz Apellániz, Nerea Huarte, Eneko Largo, and José L. Nieva. The three lives of viral fusion peptides. Chemistry and Physics of Lipids, 181:40–55, July 2014. ISSN 0009-3084. doi: 10.1016/j.chemphyslip.2014.03.003.

20. Mingfei Zhao, Laura J.S. Lopes, Harshita Sahni, Anju Yadav, Hung N. Do, Tyler Reddy, Cesar A. López, Chris Neale, and S. Gnanakaran. Insertion and Anchoring of the HIV-1 Fusion Peptide into a Complex Membrane Mimicking the Human T-Cell. The Journal of Physical Chemistry B, 128(51):12710–12727, December 2024. ISSN 1520-6106, 1520-5207. doi: 10.1021/acs.jpcb.4c05018.

21. B. Scott Perrin, Alexander J. Sodt, Myriam Cotten, and Richard W. Pastor. The Curvature Induction of Surface-Bound Antimicrobial Peptides Piscidin 1 and Piscidin 3 Varies With Lipid Chain Length. The Journal of membrane biology, 248(3):455–467, June 2015. ISSN 0022-2631. doi: 10.1007/s00232-014-9733-1.

22. Jason E. Donald, Yao Zhang, Giacomo Fiorin, Vincenzo Carnevale, David R. Slochower, Feng Gai, Michael L. Klein, and William F. DeGrado. Transmembrane orientation and possible role of the fusogenic peptide from parainfluenza virus 5 (PIV5) in promoting fusion. Proceedings of the National Academy of Sciences, 108(10):3958–3963, March 2011. doi: 10.1073/pnas.1019668108.

23. Hongwei Yao and Mei Hong. Conformation and Lipid Interaction of the Fusion Peptide of the Paramyxovirus PIV5 in Anionic and Negative-Curvature Membranes from Solid-State NMR. Journal of the American Chemical Society, 136(6):2611–2624, February 2014. ISSN 0002-7863, 1520-5126. doi: 10.1021/ja4121956.

24. Mariana Valério, Diogo A. Mendonça, João Morais, Carolina C. Buga, Carlos H. Cruz, Miguel A.R.B. Castanho, Manuel N. Melo, Cláudio M. Soares, Ana Salomé Veiga and Diana Lousa. Parainfluenza Fusion Peptide Promotes Membrane Fusion by Assembling into Oligomeric Porelike Structures. ACS Chemical Biology, 17(7):1831–1843, July 2022. ISSN 1554-8929. doi: 10.1021/acschembio.2c00208.

25. Sushmita Pal, Peter Pajtinka, Matti Javanainen, and Robert Vácha. Helical sensors of membrane saturation: Changes in orientation and curvature preference. Biophysical Journal, 124 (23):4087–4095, December 2025. ISSN 0006-3495, 1542-0086. doi: 10.1016/j.bpj.2025.09.042.

26. Piotr Setny, Paulina Borkowska, and Remigiusz Worch. Integrating Cryo-Electron Microscopy and Molecular Dynamics Simulations to Investigate Membrane Binding of Influenza Virus Fusion Peptides. Journal of the American Chemical Society, 147(16):13385–13395, April 2025. ISSN 0002-7863, 1520-5126. doi: 10.1021/jacs.4c18441.

27. Guillaume Drin and Bruno Antonny. Amphipathic helices and membrane curvature. FEBS Letters, 584(9):1840–1847, May 2010. ISSN 0014-5793, 1873-3468. doi: 10.1016/j.febslet.2009.10.022.

28. Tara C. Marcink, Tong Wang, Amedee des Georges, Matteo Porotto, and Anne Moscona. Human parainfluenza virus fusion complex glycoproteins imaged in action on authentic viral surfaces. PLoS Pathogens, 16(9):e1008883, September 2020. ISSN 1553-7366. doi: 10.1371/journal.ppat.1008883.

29. Matteo Porotto, Christine C. Yokoyama, Laura M. Palermo, Bruce Mungall, Mohamad Aljofan, Riccardo Cortese, Antonello Pessi, and Anne Moscona. Viral entry inhibitors targeted to the membrane site of action. Journal of Virology, 84(13):6760–6768, July 2010. ISSN 1098-5514. doi: 10.1128/JVI.00135-10.

30. Kayla C. Sapp, Andrew H. Beaven, and Alexander J. Sodt. Spatial extent of a single lipid’s influence on bilayer mechanics. Physical Review E, 103(4):042413, April 2021. doi: 10.1103/PhysRevE.103.042413.

31. Henry J. Lessen, Kayla C. Sapp, Andrew H. Beaven, Rana Ashkar, and Alexander J. Sodt. Molecular mechanisms of spontaneous curvature and softening in complex lipid bilayer mixtures. Biophysical Journal, 121(17):3188–3199, September 2022. ISSN 0006-3495, 1542-0086. doi: 10.1016/j.bpj.2022.07.036.

32. Amirali Hossein, Kayla Sapp, and Alexander Sodt. Computing the influence of lipids and lipid complexes on membrane mechanics. Methods in Enzymology, 701:515–540, 2024. ISSN 1557-7988. doi: 10.1016/bs.mie.2024.04.002.

33. Amirali Hossein and Alexander J. Sodt. MembraneAnalysis.jl: A Julia package for analyzing molecular dynamics simulations of lipid membranes. Journal of Open Source Software, 8 (87):5380, July 2023. ISSN 2475-9066. doi: 10.21105/joss.05380.

34. Sunhwan Jo, Taehoon Kim, Vidyashankara G. Iyer, and Wonpil Im. CHARMM-GUI: A web-based graphical user interface for CHARMM. Journal of Computational Chemistry, 29(11):1859–1865, 2008. ISSN 1096-987X. doi: 10.1002/jcc.20945. _eprint: https://onlinelibrary.wiley.com/doi/pdf/10.1002/jcc.20945.

35. James C. Phillips, David J. Hardy, Julio D. C. Maia, John E. Stone, João V. Ribeiro, Rafael C. Bernardi, Ronak Buch, Giacomo Fiorin, Jérôme Hénin, Wei Jiang, Ryan McGreevy, Marcelo C. R. Melo, Brian K. Radak, Robert D. Skeel, Abhishek Singharoy, Yi Wang, Benoît Roux, Aleksei Aksimentiev, Zaida Luthey-Schulten, Laxmikant V. Kalé, Klaus Schulten, Christophe Chipot, and Emad Tajkhorshid. Scalable molecular dynamics on CPU and GPU architectures with NAMD. The Journal of Chemical Physics, 153(4):044130, July 2020. ISSN 0021-9606. doi: 10.1063/5.0014475.

36. Jing Huang, Sarah Rauscher, Grzegorz Nawrocki, Ting Ran, Michael Feig, Bert L. de Groot, Helmut Grubmüller, and Alexander D. MacKerell. CHARMM36m: an improved force field for folded and intrinsically disordered proteins. Nature Methods, 14(1):71–73, January 2017. ISSN 1548-7105. doi: 10.1038/nmeth.4067.

37. Giacomo Fiorin, Michael L. Klein, and Jérôme Hénin. Using collective variables to drive molecular dynamics simulations. Molecular Physics, 111(22-23):3345–3362, December 2013. ISSN 0026-8976. doi: 10.1080/00268976.2013.813594. _eprint: https://doi.org/10.1080/00268976.2013.813594.

38. Sho Iketani, Ryan C. Shean, Marion Ferren, Negar Makhsous, Dolly B. Aquino, Amedee des Georges, Bert Rima, Cyrille Mathieu, Matteo Porotto, Anne Moscona, and Alexander L. Greninger. Viral Entry Properties Required for Fitness in Humans Are Lost through Rapid Genomic Change during Viral Isolation. mBio, 9(4):e00898–18, July 2018. ISSN 2150-7511. doi: 10.1128/mBio.00898-18.

39. Liqun Zhang, Alexander Bukreyev, Catherine I. Thompson, Brandy Watson, Mark E. Peeples, Peter L. Collins, and Raymond J. Pickles. Infection of ciliated cells by human parainfluenza virus type 3 in an in vitro model of human airway epithelium. Journal of Virology, 79(2): 1113–1124, January 2005. ISSN 0022-538X. doi: 10.1128/JVI.79.2.1113-1124.2005.

40. Laura M. Palermo, Manik Uppal, Lucy Skrabanek, Paul Zumbo, Soren Germer, Nora C. Toussaint, Bert K. Rima, Devra Huey, Stefan Niewiesk, Matteo Porotto, and Anne Moscona. Features of Circulating Parainfluenza Virus Required for Growth in Human Airway. mBio, 7 (2):e00235, March 2016. ISSN 2150-7511. doi: 10.1128/mBio.00235-16.

41. Samantha G. Palmer, Matteo Porotto, Laura M. Palermo, Luis F. Cunha, Olga Greengard, and Anne Moscona. Adaptation of human parainfluenza virus to airway epithelium reveals fusion properties required for growth in host tissue. mBio, 3(3):e00137–12, 2012. ISSN 2150-7511. doi: 10.1128/mBio.00137-12.

42. Alexander L. Greninger, Danielle M. Zerr, Xuan Qin, Amanda L. Adler, Reigran Sampoleo, Jane M. Kuypers, Janet A. Englund, and Keith R. Jerome. Rapid Metagenomic Next-Generation Sequencing during an Investigation of Hospital-Acquired Human Parainfluenza Virus 3 Infections. Journal of Clinical Microbiology, 55(1):177–182, January 2017. ISSN 1098-660X. doi: 10.1128/JCM.01881-16.

43. David N. Mastronarde. Automated electron microscope tomography using robust prediction of specimen movements. Journal of Structural Biology, 152(1):36–51, October 2005. ISSN 1047-8477. doi: 10.1016/j.jsb.2005.07.007.

44. Dimitry Tegunov and Patrick Cramer. Real-time cryo-electron microscopy data preprocessing with Warp. Nature Methods, 16(11):1146–1152, November 2019. ISSN 1548-7105. doi: 10.1038/s41592-019-0580-y.

45. Shawn Zheng, Georg Wolff, Garrett Greenan, Zhen Chen, Frank G. A. Faas, Montserrat Bárcena, Abraham J. Koster, Yifan Cheng, and David A. Agard. AreTomo: An integrated software package for automated marker-free, motion-corrected cryo-electron tomographic alignment and reconstruction. Journal of Structural Biology: X, 6:100068, 2022. ISSN 2590-1524. doi: 10.1016/j.yjsbx.2022.100068.

46. Dimitry Tegunov, Liang Xue, Christian Dienemann, Patrick Cramer, and Julia Mahamid. Multiparticle cryo-EM refinement with M visualizes ribosome-antibiotic complex at 3.5{\AA} in cells. Nature Methods, 18(2):186–193, February 2021. ISSN 1548-7105. doi: 10.1038/s41592-020-01054-7.

47. Josh Abramson, Jonas Adler, Jack Dunger, Richard Evans, Tim Green, Alexander Pritzel, Olaf Ronneberger, Lindsay Willmore, Andrew J. Ballard, Joshua Bambrick, Sebastian W. Bodenstein, David A. Evans, Chia-Chun Hung, Michael O’Neill, David Reiman, Kathryn Tunyasuvunakool, Zachary Wu, Akvilė Žemgulytė, Eirini Arvaniti, Charles Beattie, Ottavia Bertolli, Alex Bridgland, Alexey Cherepanov, Miles Congreve, Alexander I. Cowen-Rivers, Andrew Cowie, Michael Figurnov, Fabian B. Fuchs, Hannah Gladman, Rishub Jain, Yousuf A. Khan, Caroline M. R. Low, Kuba Perlin, Anna Potapenko, Pascal Savy, Sukhdeep Singh, Adrian Stecula, Ashok Thillaisundaram, Catherine Tong, Sergei Yakneen, Ellen D. Zhong, Michal Zielinski, Augustin Žídek, Victor Bapst, Pushmeet Kohli, Max Jaderberg, Demis Hassabis, and John M. Jumper. Accurate structure prediction of biomolecular interactions with AlphaFold 3. Nature, 630(8016):493–500, June 2024. ISSN 1476-4687. doi: 10.1038/s41586-024-07487-w.

48. Michael M. Kozlov and Justin W. Taraska. Generation of nanoscopic membrane curvature for membrane trafficking. Nature Reviews Molecular Cell Biology, 24(1):63–78, January 2023. ISSN 1471-0072, 1471-0080. doi: 10.1038/s41580-022-00511-9.

49. Yonathan Kozlovsky, Joshua Zimmerberg, and Michael M. Kozlov. Orientation and interaction of oblique cylindrical inclusions embedded in a lipid monolayer: a theoretical model for viral fusion peptides. Biophysical Journal, 87(2):999–1012, August 2004. ISSN 0006-3495. doi: 10.1529/biophysj.104.041467.

50. Joshua Zimmerberg and Michael M. Kozlov. How proteins produce cellular membrane curvature. Nature Reviews Molecular Cell Biology, 7(1):9–19, January 2006. ISSN 1471-0072, 1471-0080. doi: 10.1038/nrm1784.

51. Ruthellen H. Anderson, Kem A. Sochacki, Harika Vuppula, Brandon L. Scott, Elizabeth M. Bailey, Maycie M. Schultz, Jason G. Kerkvliet, Justin W. Taraska, Adam D. Hoppe, and Kevin R. Francis. Sterols lower energetic barriers of membrane bending and fission necessary for efficient clathrin-mediated endocytosis. Cell Reports, 37(7):110008, November 2021. ISSN 22111247. doi: 10.1016/j.celrep.2021.110008.

52. Remigiusz Worch, Anita Dudek, Paulina Borkowska, and Piotr Setny. Transient Excursions to Membrane Core as Determinants of Influenza Virus Fusion Peptide Activity. International Journal of Molecular Sciences, 22(10):5301, January 2021. ISSN 1422-0067. doi: 10.3390/ijms22105301.

53. Jatin Soni and Taraknath Mandal. Investigating the Impact of Cholesterol on GP41-Driven HIV Membrane Fusion. Journal of Chemical Theory and Computation, 21(18):9164–9174, September 2025. ISSN 1549-9618, 1549-9626. doi: 10.1021/acs.jctc.5c00962.

54. Mariana Valério, Carolina C. Buga, Diogo A. Mendonça, Miguel A. R. B. Castanho, Manuel N. Melo, Cláudio M. Soares, Diana Lousa, and Ana Salomé Veiga. Unravelling the role of key amino acid residues of the parainfluenza fusion peptide in membrane fusion. RSC Chemical Biology, 6(7):1100–1114, 2025. ISSN 2633-0679. doi: 10.1039/D5CB00058K.

55. Kyle Stearns, Tara Marcink, Emily Pawlack, Elizabeth B. Sobolik, Matteo Porotto, Alexander L. Greninger, Stefan Nieweisk, and Anne Moscona. Human parainfluenza virus 3 fusion protein cleavage: a key determinant of infection and spread. Journal of Virology, 100(6):e02126–25, June 2026. ISSN 0022-538X, 1098-5514. doi: 10.1128/jvi.02126-25.

56. Per Larsson and Peter M. Kasson. Lipid Tail Protrusion in Simulations Predicts Fusogenic Activity of Influenza Fusion Peptide Mutants and Conformational Models. PLoS Computational Biology, 9(3):e1002950, March 2013. ISSN 1553-7358. doi: 10.1371/journal.pcbi.1002950.

57. Chetan S. Poojari, Katharina C. Scherer, and Jochen S. Hub. Free energies of membrane stalk formation from a lipidomics perspective. Nature Communications, 12(1):6594, November 2021. ISSN 2041-1723. doi: 10.1038/s41467-021-26924-2.

58. Katharina C. Scherer, Chetan S. Poojari, and Jochen S. Hub. Transmembrane domains of fusion proteins promote stalk formation by inducing membrane disorder. Biophysical Journal, 125(5):1276–1285, March 2026. ISSN 00063495. doi: 10.1016/j.bpj.2026.01.032.

59. D. P. Siegel and M. M. Kozlov. The Gaussian Curvature Elastic Modulus of N-Monomethylated Dioleoylphosphatidylethanolamine: Relevance to Membrane Fusion and Lipid Phase Behavior. Biophysical Journal, 87(1):366–374, July 2004. ISSN 0006-3495. doi: 10.1529/biophysj.104.040782.

60. Vladimir I. Razinkov, Grigory B. Melikyan, Richard M. Epand, Raquel F. Epand, and Fredric S. Cohen. Effects of Spontaneous Bilayer Curvature on Influenza Virus–mediated Fusion Pores. The Journal of General Physiology, 112(4):409–422, October 1998. ISSN 0022-1295. doi: 10.1085/jgp.112.4.409.

61. L. Chernomordik, A. Chanturiya, J. Green, and J. Zimmerberg. The hemifusion intermediate and its conversion to complete fusion: regulation by membrane composition. Biophysical Journal, 69(3):922–929, September 1995. ISSN 0006-3495. doi: 10.1016/S0006-3495(95)79966-0.

62. Leonid V. Chernomordik, Eugenia Leikina, Vadim Frolov, Peter Bronk, and Joshua Zimmerberg. An Early Stage of Membrane Fusion Mediated by the Low pH Conformation of Influenza Hemagglutinin Depends upon Membrane Lipids. Journal of Cell Biology, 136(1):81–93, January 1997. ISSN 0021-9525. doi: 10.1083/jcb.136.1.81.

63. Andrew H. Beaven, Kayla Sapp, and Alexander J. Sodt. Simulated dynamic cholesterol redistribution favors membrane fusion pore constriction. Biophysical Journal, 122(11):2162–2175, June 2023. ISSN 00063495. doi: 10.1016/j.bpj.2022.12.024.

64. Leonid V. Chernomordik and Michael M. Kozlov. Mechanics of membrane fusion. Nature Structural & Molecular Biology, 15(7):675–683, July 2008. ISSN 1545-9985. doi: 10.1038/nsmb.1455.

