## Supplementary Information for "Fusion Peptide Insertion Configuration Determines the Mechanism of Membrane Curvature Induction in Human Parainfluenza Virus Type 3"

### Data Analysis

#### Fusion Peptide Residue Height

FP residue heights were calculated from all-atom MD trajectories using MDAnalysis [1, 2]. For each frame within the analysis window, the z-coordinate of the $C_\alpha$ atom of each FP residue (residues 1-32) was extracted for all three helices. Residue height was defined relative to the simulation coordinate system, with the bilayer midplane located at approximately $z = 0$ Å. For each replicate, residue heights were averaged over time to generate residue-specific height profiles. Residue heights were subsequently averaged across replicate simulations using frame-weighted averaging. Variability was quantified using pooled standard deviations that account for both fluctuations within individual simulations and differences between replicate averages. The average positions of the membrane neutral surface and upper-leaflet POPC phosphate groups were calculated and used as membrane reference planes. Residue height profiles are reported as pooled means, and shaded regions represent pooled standard deviations.

#### Lipid Traces

Lipid traces were computed similarly to Ref. [3] using code available at github.com/alexsodt/LipidTrace (commit f084afc). Briefly, the distance from the center-of-geometry of a lipid to the center-of-geometry of a peptide segment is used to assign the lipid to a histogram of *lipid distances*. The histogram is built from the entire trajectory. The histogram is then parsed beginning with lipids closest to the protein. Lipids are averaged until the total number meets a target threshold, giving a set of lipids whose lipid distances are all within a limited range, indexed by $i$. For each atom of the lipid, the average $r_i$ and $z_i$ position is computed, and plotted, with line connections between bonded atoms, with branches ignored (for example, hydrogens, methyl, and carbonyl groups). By averaging over lipids selected by the lipid-distance histogram, each trace reflects an average of the atoms of a *whole* lipid. For radial averaging (for TM helices), a weight of $r_{xy}^{-1}$ is added for each atom, to correct for the geometric/entropic factor that favors radially farther atoms. For averaging perpendicular to surface-adsorbed helices, there is no geometric bias so the factor is not included.

#### Radial Distribution Function

Radial distribution functions (RDF's) were computed with a simple in-house program (`http://github.com/alexsodt/gofr`, commit e266413) that produces equivalent distributions as other standard packages. Distribution functions were resolved at a bon width of 0.1 Å.

#### Visualization

Molecular renderings were generated using Visual Molecular Dynamics (VMD) version 2.0 [4]. Representative snapshots from replicate 1 production trajectories were selected to illustrate FP orientation within the membrane. For visualizations focused on local FP-lipid interactions, only the FP and neighboring lipid molecules were displayed to highlight lipid orientations arising from these interactions.

#### Membrane Curvature Analysis of Cryo-Electron Tomography Slices

Cryo-ET tomograms of activated HPIV3 bound to host erythrocyte (RBC) membrane vesicles were analyzed for regions of virus-host interactions. 2D slices from these regions were analyzed to quantify localized host-cell membrane deformation at virus-cell interfaces. Host-cell membrane segments interacting with HPIV3 were manually annotated to define regions of interest for automated analysis. These annotations were used only to identify the region of virus-host interactions. All quantitative measurements were performed directly from the cryo-ET 2D slices.

Membrane centerlines were reconstructed using a custom Python pipeline built on scikit-image that combined algorithms for multiscale pixel edge detection [5] with shortest-path optimization to identify the outer edge of the membrane within each annotated region. The resulting membrane traces were smoothed using quintic spline interpolation to generate continuous membrane contours. Local membrane curvature was calculated from the spline derivatives along the membrane trace. To identify the site of HPIV3-induced membrane deformation, the point of closest proximity between the host-cell and viral membranes was determined, and the local curvature at this position ($J_{\text{local}}$) was calculated as the mean curvature within a ±20 nm window centered on the contact point. Averaging over a finite arc length reduced sensitivity to local image noise while preserving localized membrane geometry.

Because host-cell membrane vesicles are pleomorphic and enclose different volumes, the baseline curvature of each membrane segment was estimated independently by fitting an algebraic least-squares circle to the reconstructed membrane contour. The resulting radius ($R_{\text{circle}}$) was used to calculate the expected curvature of the undeformed membrane ($J_{\text{circle}}$). Local membrane deformation associated with viral attachment was quantified as the excess curvature at the contact site ($\Delta J = J_{\text{local}} - J_{\text{circle}}$), allowing curvature measurements to be compared across membrane vesicles of different sizes. Positive or negative $\Delta J$ values indicate that the membrane is curving towards the virus or away from it, respectively, at the virus-cell interface compared to the calculated optimal spherical radii.

All reconstructed membrane traces were visually inspected following automated analysis, and traces that did not accurately follow the membrane density were excluded before downstream analysis. The complete analysis pipeline was implemented in Python using NumPy [6], SciPy [7], scikit-image [8], OpenCV, and Matplotlib [9].

### Statistical Methods

Molecular dynamics trajectories were run in triplicate for each combination of system (WT 3TM, F1A/F2A, NNT, WT 3SP, WT 1TM, WT 1SP) and cholesterol concentration (0%, 10%, 30%; N = 42 replicates,with 14 of the 18 possible system–cholesterol combinations represented), yielding per-replicate $\langle J \rangle A_{\text{bilayer}}$ computed over 231-901 frames per trajectory. Each trajectory was analyzed by two-way ANOVA with system, cholesterol, and their interaction as fixed factors, using cell-means general linear models and Type I (sequential) sums of squares. Residual normality and variance homogeneity were assessed via Shapiro-Wilk and Jarque-Bera tests and Levene's test. Significant omnibus effects were followed by Tukey's Honestly Significant Difference (HSD) post-hoc comparisons using the pooled residual mean square error. As a sensitivity analysis, the ANOVA and its post-hoc comparisons were repeated with replicates weighted by trajectory length (number of frames) via weighted least squares (Tables II and III). All analyses were performed in custom JavaScript and MATLAB 2025b implementations (numerical linear algebra via mathjs and MATLAB's built-in matrix operations; distributional and hypothesis-testing functions via MATLAB's Statistics and Machine Learning Toolbox, supplemented by custom implementations of the Shapiro-Wilk test and studentized range distribution where native functions were unavailable) Significance was assessed at $\alpha = 0.05$.

The N = 42 replicates were annotated with two additional structural descriptors of each system, Peptide Number (Trimer, Monomer) and Geometry (Trans, Surface). Because each System maps to exactly one Peptide Number × Geometry combination, these three factors (System, Peptide Number, and Geometry) are structurally confounded. System's effect was therefore decomposed hierarchically into Peptide Number, Geometry, their interaction, and a System-within-cell residual term (Table IV). Because the System × Cholesterol interaction was significant in the two-way ANOVA, Peptide Number and Geometry were additionally re-examined separately at 0% and 30% cholesterol (a "simple effects" follow-up).

Cholesterol concentration was additionally modeled as a continuous predictor via simple linear regression ($\langle J \rangle A_{\text{bilayer}} \sim \beta_0 + \beta_1 \times \text{Cholesterol}$) and fit separately for each system. Systems were classified by the sign and significance of their slope: three systems (WT 3TM, F1A/F2A, NNT) showed a significant negative slope and were carried forward for zero-crossing estimation ($x_0 = -\beta_0/\beta_1$, with 95% CI via Fieller's theorem). The remaining three systems (WT 3SP, WT 1TM, WT 1SP) were evaluated only for slope significance. The categorical System × Cholesterol interaction (two-way ANOVA) was independently confirmed using cholesterol as a continuous covariate via a heterogeneous-slopes nested F-test.

Bootstrap resampling (10,000 iterations) with bias-corrected and accelerated (BCa) 95% confidence intervals was used to estimate the mean, variance, and percentage of positive/negative values for each dataset (Clinical Isolate, N = 604; HN-P111S Variant, N = 121), and a two-sample bootstrap hypothesis test was performed to assess evidence against the null hypotheses of equal mean and equal variance between groups.

### I. SUPPLEMENTAL TABLE 1

**Table S1. Summary of molecular dynamics simulation systems and restraint conditions.**
Simulation systems are listed according to membrane cholesterol content, initial protein orientation, amino acid substitutions, simulation time for each replicate, and the collective-variable restraint applied.

| Chol (%) | Orientation | Amino acid substitutions | Simulation time per replicate ($\mu$s) | Restraint |
|---|---|---|---|---|
| **Trimer Systems** | | | | |
| 0 | TM | WT | 4.00, 3.85, 3.80 | $\alpha$-helix |
| 10 | TM | WT | 5.00, 5.00, 5.00 | $\alpha$-helix |
| 30 | TM | WT | 5.00, 5.00, 5.00 | $\alpha$-helix |
| 0 | TM | F1A/F2A | 1.95, 1.65, 2.20 | $\alpha$-helix |
| 30 | TM | F1A/F2A | 4.40, 4.15, 4.00 | $\alpha$-helix |
| 0 | TM | NNT | 2.15, 2.15, 1.95 | $\alpha$-helix |
| 30 | TM | NNT | 3.95, 3.90, 4.20 | $\alpha$-helix |
| 0 | SP | WT | 4.00, 4.55, 4.30 | RMSD |
| 10 | SP | WT | 4.25, 5.00, 4.10 | RMSD |
| 30 | SP | WT | 5.00, 5.00, 4.05 | RMSD |
| **Monomer Systems** | | | | |
| 10 | TM | WT | 5.00, 5.00, 5.00 | $\alpha$-helix |
| 30 | TM | WT | 5.00, 5.00, 5.00 | $\alpha$-helix |
| 10 | SP | WT | 5.00, 5.00, 5.00 | $\alpha$-helix |
| 30 | SP | WT | 5.00, 5.00, 5.00 | $\alpha$-helix |

**Abbreviations:** TM, transmembrane; SP, surface-parallel; WT, wild type; F1A/F2A, double phenylalanine-to-alanine substitution mutant. NNT, neutral N-terminus. Trimer TM and all monomer simulations were restrained using the Colvars $\alpha$-helicity collective variable with a harmonic force constant of 3.5 kcal mol$^{-1}$. Trimer SP simulations were restrained using an RMSD collective variable with a harmonic force constant of 3.5 kcal mol$^{-1}$.

### II. SUPPLEMENTAL TABLE 2

**Table S2. Frames-weighted two-way ANOVA of $\langle J \rangle_A$ by system and cholesterol concentration.** As a sensitivity analysis, the two-way ANOVA was performed using weighted least squares, with each replicate weighted by its number of analyzed trajectory frames ($N_{\text{frames}}$), such that longer trajectories contributed more strongly to the analysis. System, cholesterol concentration, and their interaction were included as fixed effects.

| Source | df | SS | MS | F | p |
|---|---|---|---|---|---|
| System | 5 | 581,375.9 | 116,275.2 | 7.49 | 0.00014 |
| Cholesterol | 2 | 122,055.1 | 61,027.6 | 3.93 | 0.031 |
| System × Cholesterol | 6 | 324,460.1 | 54,076.7 | 3.49 | 0.011 |
| Residual | 28 | 434,511.2 | 15,518.3 | | |

**Abbreviations:** df, degrees of freedom. SS, sum of squares. MS, mean square. F, F-statistic. p, p-value.
**Number of frames:** $N_{\text{frames}} = \left\lfloor \frac{(\text{time}_{\text{ns}})(\frac{1\text{frame}}{ns})-500}{5} \right\rfloor$

### III. SUPPLEMENTAL TABLE 3

**Table S3. Significant Tukey HSD post-hoc comparisons following the frames-weighted ANOVA.**
Tukey honestly significant difference (HSD) post-hoc comparisons were performed following the frames-weighted two-way ANOVA. Only comparisons that were statistically significant ($p < 0.05$) are shown. Differences are reported as the first listed group minus the second listed group.

| Comparison | Difference | p |
|---|---|---|
| 3-SP vs 1-TM | +10.16 | 0.0013 |
| 3-TM vs 1-TM | +9.63 | 0.0022 |
| NNT vs 3-SP | -10.10 | 0.0095 |
| 3-TM vs NNT | +9.57 | 0.0146 |
| F1A/F2A vs 3-SP | -8.53 | 0.038 |
| 0% vs 30% Cholesterol | +7.51 | 0.0013 |

**Abbreviations:** 1-TM, WT transmembrane monomer; 3-TM, WT transmembrane trimer; 3-SP, WT surface-parallel trimer; F1A/F2A, double phenylalanine-to-alanine substitution mutant; NNT, neutral N-terminus.

### IV. SUPPLEMENTAL TABLE 4

**Table S4. Nested ANOVA decomposition of system effects on $\langle J \rangle A_{\text{bilayer}}$.**

The System effect from the two-way ANOVA was decomposed into peptide number (monomer or trimer), geometry (transmembrane or surface-parallel), their interaction, and residual system variation within peptide number × geometry. Cholesterol concentration and the System × Cholesterol interaction were retained from the original model. Type I (sequential) sums of squares are reported.

| Source | df | SS | MS | F | p |
|---|---|---|---|---|---|
| Peptide Number | 1 | 169.27 | 169.27 | 9.19 | 0.0052 |
| Geometry | 1 | 206.65 | 206.65 | 11.21 | 0.0023 |
| Peptide × Geometry | 1 | 3.88 | 3.88 | 0.21 | 0.650 |
| System (with Peptide × Geometry) | 2 | 256.69 | 128.34 | 6.96 | 0.0035 |
| Cholesterol | 2 | 279.72 | 139.86 | 7.59 | 0.0023 |
| System × Cholesterol | 6 | 476.84 | 79.47 | 4.31 | 0.0034 |
| Residual | 28 | 516.00 | 18.43 | | |

**Abbreviations:** df, degrees of freedom; SS, sum of squares; MS, mean square; F, F-statistic; p, p-value; TM, transmembrane; SP, surface-parallel.

### V. SUPPLEMENTAL TABLE 5

**Table S5. Summary statistics and bootstrap comparisons of Clinical Isolate and HN-P111S variant cryo-ET datasets.**

Summary statistics were calculated for the Clinical Isolate and HN-P111S variant datasets. Mean and variance are reported for each dataset together with bias-corrected and accelerated (BCa) 95% bootstrap confidence intervals. Between-group differences in the mean and variance were calculated as Clinical Isolate − HN-P111S variant and evaluated based on whether the corresponding BCa 95% confidence interval included zero. The proportions of observations greater than and less than zero are also reported with BCa 95% bootstrap confidence intervals.

| Measure | Dataset / Comparison | N | Estimate | BCa 95% CI | Inference |
|---|---|---|---|---|---|
| **A. Distribution Summary** | | | | | |
| Mean | Clinical Isolate | 604 | 1.8007 | [1.3927, 2.2325] | – |
| Mean | HN-P111S variant | 121 | −0.9628 | [−2.0405, −0.2605] | – |
| Variance | Clinical Isolate | 604 | 27.3679 | [22.3220, 36.0305] | – |
| Variance | HN-P111S variant | 121 | 23.3016 | [9.9451, 55.4659] | – |
| **B. Between-Group Comparison** | | | | | |
| Mean difference | Clinical Isolate − HN-P111S variant | – | 2.7635 | [1.9479, 3.8884] | CI excludes 0 |
| Variance difference | Clinical Isolate − HN-P111S variant | – | 4.0663 | [−26.6614, 18.1163] | CI includes 0 |
| **C. Values Relative to Zero** | | | | | |
| Values > 0 | Clinical Isolate | 604 | 63.74% | [59.77%, 67.38%] | – |
| Values > 0 | HN-P111S variant | 121 | 44.63% | [35.54%, 52.89%] | – |
| Values < 0 | Clinical Isolate | 604 | 36.26% | [32.45%, 40.07%] | – |
| Values < 0 | HN-P111S variant | 121 | 55.37% | [46.28%, 63.64%] | – |

**Abbreviations:** N, number of observations; BCa, bias-corrected and accelerated; CI, confidence interval.
Between-group differences were calculated as Clinical Isolate − HN-P111S variant. For between-group comparisons, a BCa 95% CI excluding zero indicates evidence of a difference between groups.

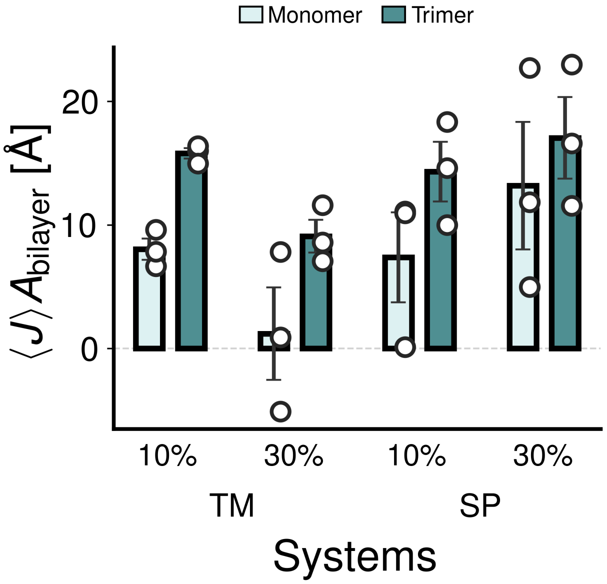


FIG. S1. Comparison of membrane curvature $\langle J \rangle A_{\mathrm{bilayer}}$ generated by HPIV3 FP monomer and trimer systems in the surface-parallel (SP) and transmembrane (TM) configurations for membranes containing 10% and 30% cholesterol. Bars indicate the mean value averaged over time and replicates, circles denote individual replicate means, and error bars represent the standard error of the mean.

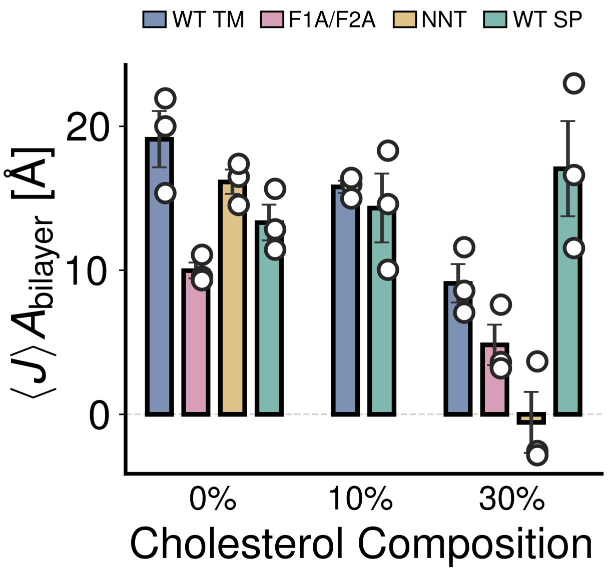


FIG. S2. Comparison of membrane curvature $\langle J \rangle A_{\text{bilayer}}$ generated by WT HPIV3 FP trimers in the surface-parallel (SP) and transmembrane (TM) configurations at 0%, 10%, and 30% cholesterol. Curvature generated by the F1A/F2A and neutral N-terminus (NNT) transmembrane systems is shown for comparison. Bars indicate the mean value averaged over time and replicates, circles denote individual replicate means, and error bars represent the standard error of the mean.

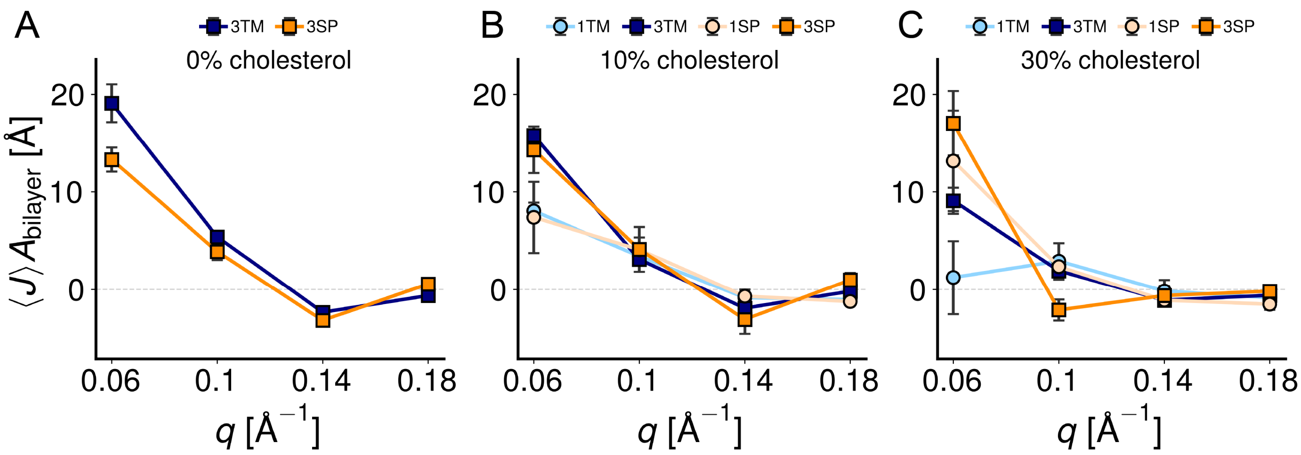


FIG. S3. Spontaneous curvature spectra sampled from simulations of the HPIV3 FP trimer and monomer in different configurations and cholesterol concentrations. FP transmembrane configuration is shown in blue. Surface-parallel configuration is depicted in orange. Trimers are represented with a square marker. Monomers are represented with a circular marker. Each data point was the averaged in time in each mode. Error bars represent the standard error of the mean.

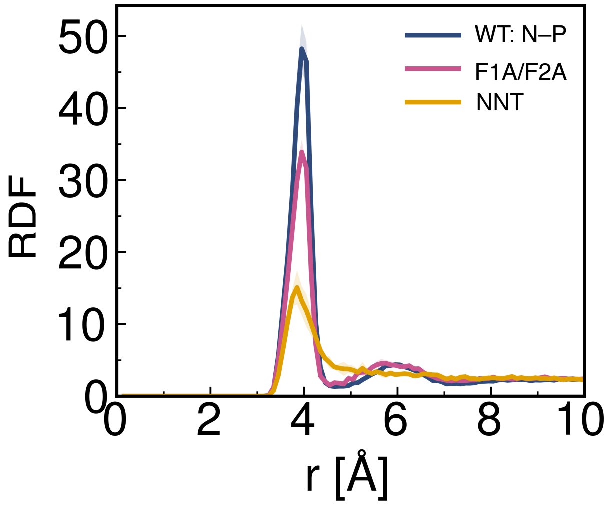


FIG. S4. Radial distribution function, RDF, calculated between the backbone N atom of the first FP residue and lipid phosphate (P) atoms for the 0% cholesterol systems, showing a progressive reduction in the first peak amplitude from the WT to the F1A/F2A and NNT systems.

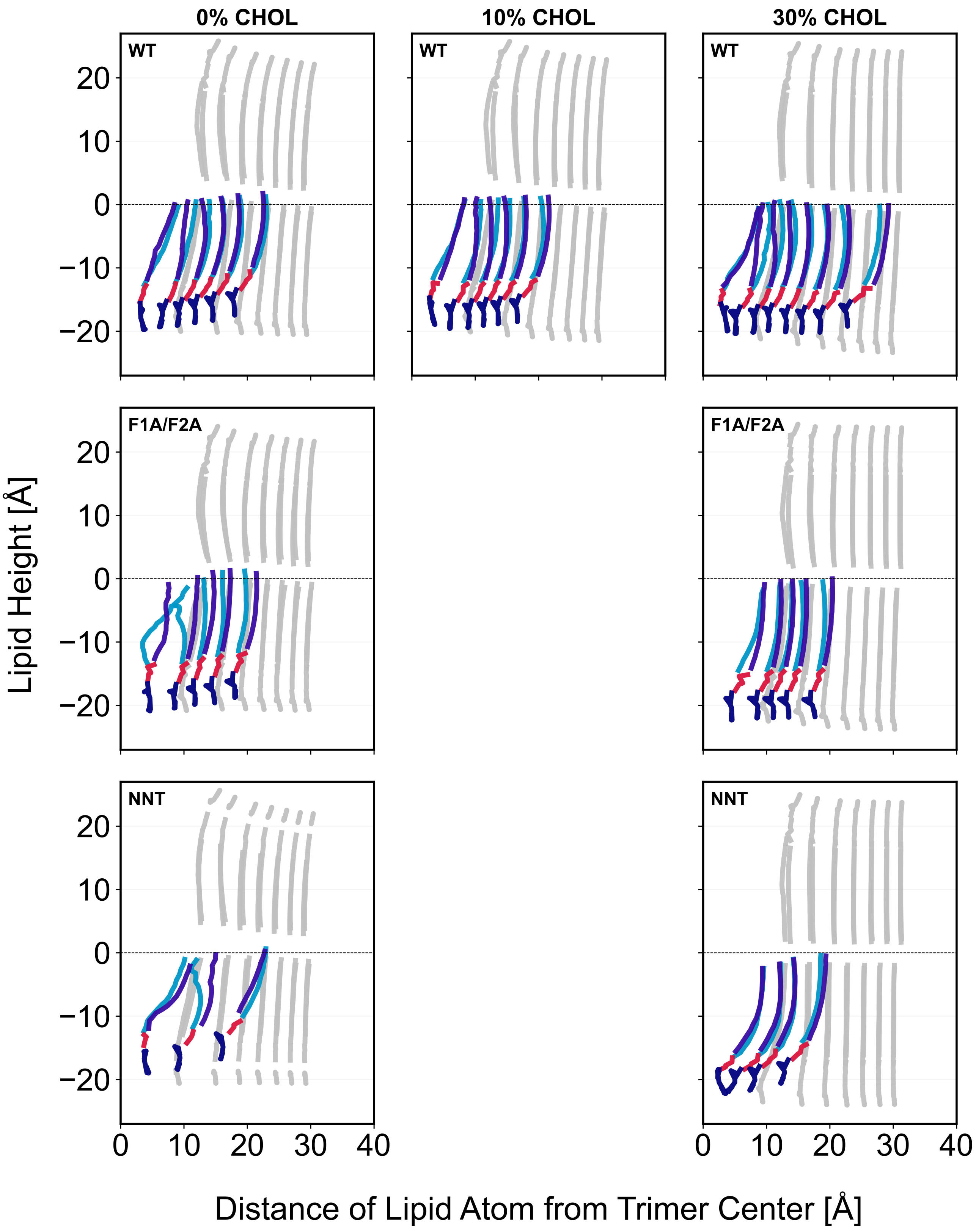


FIG. S5. Local lipid organization around the HPIV3 FP N-termini across cholesterol concentrations and FP variants. Lipid height is shown as a function of radial distance from the trimer center for WT, F1A/F2A, and neutral N-terminus (NNT) systems at the indicated cholesterol concentrations. Colored traces represent individual lipids located within 4.75 Å of an FP N-terminus. Gray traces show the average lipid height as a function of distance from the trimer center, calculated by averaging lipids within concentric radial bins around the trimer. The horizontal dashed line indicates a lipid height of 0 Å. Conditions that were not simulated are left blank.

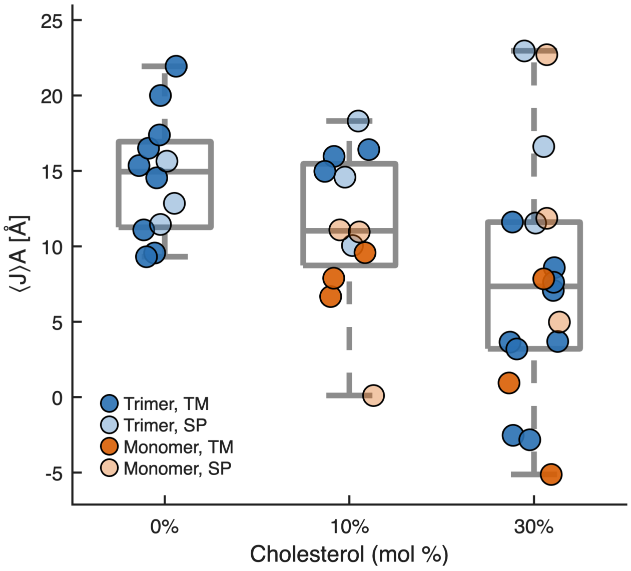


FIG. S6. $\langle J \rangle A$ declines with increasing cholesterol. Boxplots of $\langle J \rangle A$ at 0%, 10%, and 30% cholesterol (points jittered, n = 12-18 per level, pooled across all six systems). Cholesterol concentration was a significant predictor of $\langle J \rangle A$ (two-way ANOVA, partial $\eta^2 = 0.351$ , F(2,28) = 7.59, p = 0.0023); the 0%-vs-30% contrast was the strongest pairwise driver (Tukey HSD, p = 0.0003, Cohen's d = 1.08).

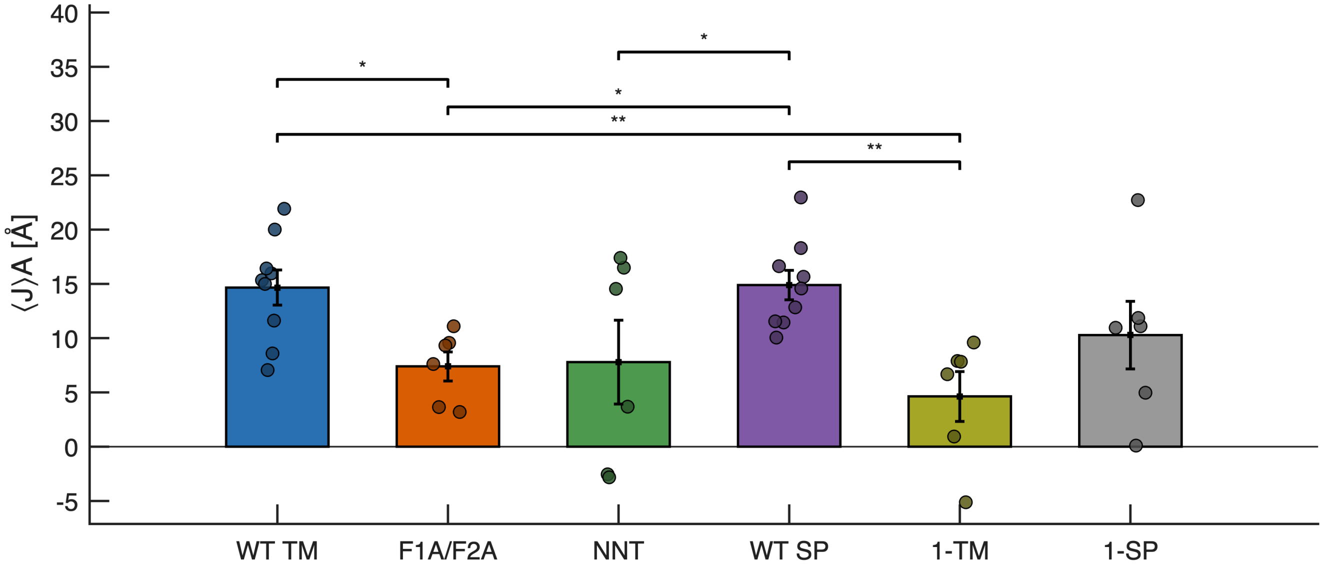


FIG. S7. $\langle J \rangle A$ varies significantly by system. Bar plot of $\langle J \rangle A$ by system (mean ± SEM, individual replicates overlaid), pooled across cholesterol levels. System was a significant, large-effect predictor of ⟨J⟩A (partial $\eta^2$ = 0.552, F(5,28) = 6.91, p = 0.00026), driven primarily by 1-TM being distinctly lower than WT TM and WT SP; five pairwise contrasts reached significance by Tukey HSD, all corresponding to large effect sizes (Cohen's d = 1.06-2.17).

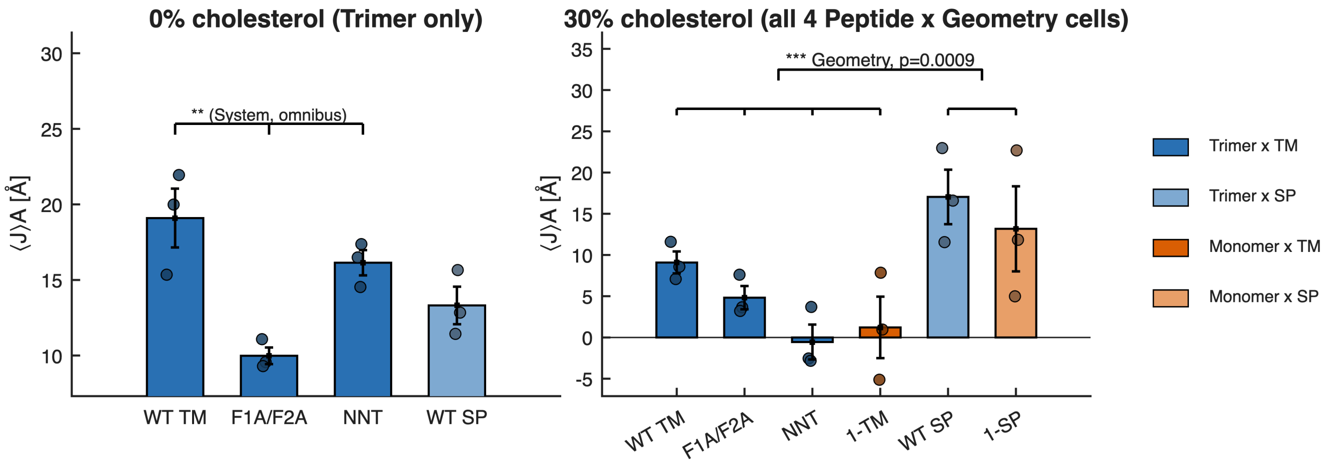


FIG. S8. Cholesterol-stratified analysis of system and geometry effects on membrane curvature. $\langle J \rangle A_{\mathrm{bilayer}}$ at 0% (left) and 30% (right) cholesterol. At 0%, only trimer systems were available; WT TM, F1A/F2A, and NNT (all transmembrane) differed significantly by omnibus test (p = 0.0027), while WT SP is shown for reference. At 30%, all four peptide-number $\times$ geometry combinations were available, allowing peptide number and geometry to be evaluated independently; geometry remained significant (p = 0.0009). Bars show group means $\pm$ SEM; circles show individual replicates.

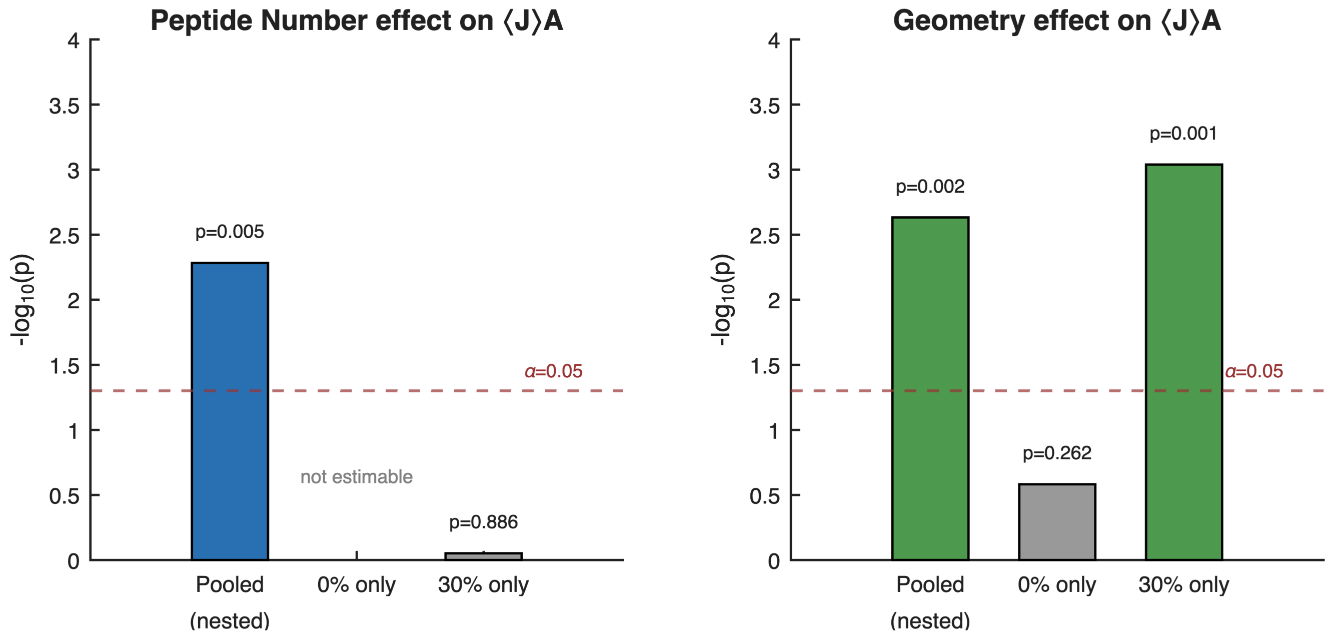


FIG. S9. Peptide Number's pooled significance is an artifact of confounding with cholesterol; Geometry's is not. Bar height represents $-\log_{10}(p)$ for the effect of Peptide Number (left) and Geometry (right) on $\langle J \rangle A$, compared across three analyses: the pooled nested model, and the restricted-level models holding cholesterol fixed at 0% or 30%. The dashed line marks the $\alpha = 0.05$ significance threshold. Peptide Number is significant in the pooled model (p=0.005) but not estimable at 0% (no Monomer systems tested at that concentration) and falls short of significance at 30% (p=0.886) — the only level at which it can be directly tested — indicating the pooled effect substantially reflects confounding with cholesterol rather than a genuine, condition-independent structural effect. Geometry, by contrast, remains significant in the pooled model (p=0.002) and at 30% (p=0.001), and while not significant at 0% alone (p=0.262, likely reflecting the smaller n=12 at that level), its effect direction is consistent across both tested levels, supporting Geometry as the more robust structural predictor of $\langle J \rangle A$.

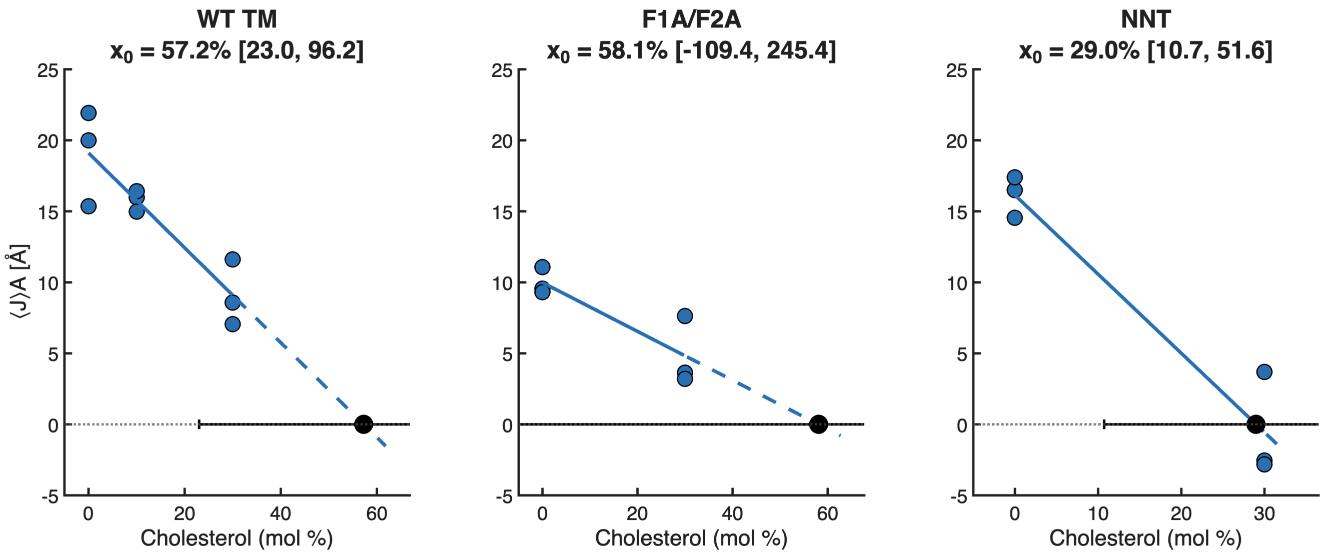


FIG. S10. Zero-crossing estimation for systems with a significant declining $\langle J\rangle A$- cholesterol slope. Per-system linear regression of ⟨J⟩A against continuous cholesterol concentration for WT TM, F1A/F2A, and NNT (all $p \leq 0.03$ for slope, $R^2 = 0.74 - 0.93$), with the estimated cholesterol concentration at which $\langle J\rangle A$ reaches zero ($x_0$) and its Fieller 95% confidence interval. NNT provided the most precise, non-extrapolated estimate ($x_0$ = 29.0%, 95% CI 10.7-51.6%, within the tested dose range); WT TM's estimate (57.2%) required substantial extrapolation, and F1A/F2A's confidence interval (-109.4% to 245.4%) was too wide to support a precise value.

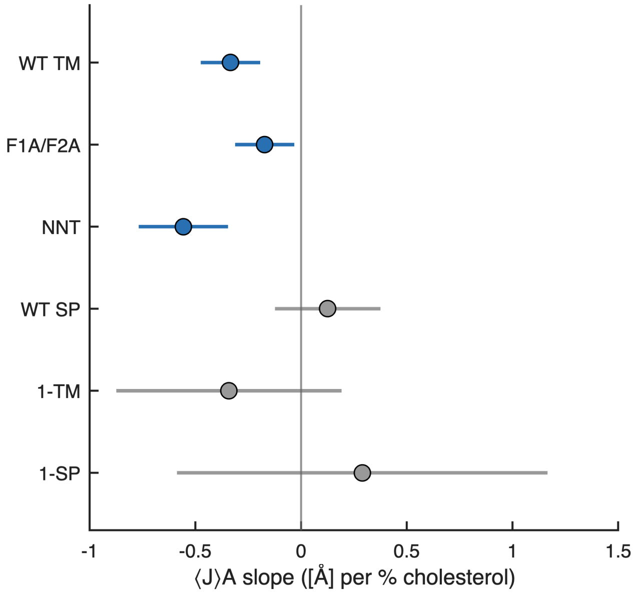


FIG. S11. Cholesterol slope heterogeneity across all six systems. Forest plot of per-system $\langle J \rangle A$-vs-cholesterol slopes with 95% confidence intervals, colored by significance. Three systems (WT TM, F1A/F2A, NNT) show significant negative slopes; the remaining three (WT SP, 1-TM, 1-SP) do not reach significance, though 1-TM's slope (-0.341) is numerically consistent with the declining systems and reflects low statistical power (n = 6) rather than a true absence of effect. This system-level heterogeneity was formally confirmed by a heterogeneous-slopes F-test (F(5,30) = 5.84, p = 0.0007)

**Expected curvature deformation over a region of a sphere**

Consider a vesicle surface parameterized in spherical coordinates. The radius has the form

$$r(\theta,\phi) = R\left[1+u(\theta,\phi)\right] \tag{S1}$$

such that the surface is

$$\mathbf{r} = r(\theta,\phi)\{\sin(\theta)\cos(\phi), \sin(\theta)\sin(\phi), \cos(\theta)\}. \tag{S2}$$

With the intention of putting a cap on the pole representing the protein's local effect on spontaneous curvature, we assume axial symmetry such that derivatives with respect to $\phi$ ($u_\phi$, $u_{\phi\phi}$) are zero.

The first fundamental form is derived from

$$E = \frac{\partial\mathbf{r}}{\partial\theta}\cdot\frac{\partial\mathbf{r}}{\partial\theta} = R^2\left((1+u)^2 + u_\theta^2\right) \tag{S3}$$

$$F = \frac{\partial\mathbf{r}}{\partial\theta}\cdot\frac{\partial\mathbf{r}}{\partial\phi} = 0 \tag{S4}$$

$$G = \frac{\partial\mathbf{r}}{\partial\phi}\cdot\frac{\partial\mathbf{r}}{\partial\phi} = R^2\sin^2(\theta)(1+u)^2 \tag{S5}$$

$$\tag{S6}$$

such that the area element (the length of the cross product of $\frac{\partial\mathbf{r}}{\partial\theta}$ with $\frac{\partial\mathbf{r}}{\partial\phi}$) is (to second order in $u$):

$$\sqrt{EG-F^2} = R^2\sin(\theta)\left(1+2u+u^2+\frac{1}{2}\left(u_\theta^2\right)\right) + \mathcal{O}\left[u^3\right]. \tag{S7}$$

This sets the area element for integration as

$$\mathrm{d}A = R^2\left(1+2u+u^2+\frac{1}{2}(u_\theta)^2\right)\mathrm{d}\Omega \tag{S8}$$

with

$$\mathrm{d}\Omega = \sin(\theta)\mathrm{d}\theta\mathrm{d}\phi \tag{S9}$$

With spontaneous curvature set for an end cap pole of a spherical particle it is not necessary to carry terms $u_\phi$ that break axial symmetry.

The second fundamental form neglecting terms $u_\phi$ and expanding to second order in $u$ is

$$L = \frac{\partial^2\mathbf{r}}{\partial\theta^2}\cdot\mathbf{n} = -R\left(1+\frac{1}{2}\left(2u+3u_\theta^2-2u_{\theta\theta}\right)\right) \tag{S10}$$

$$M = \frac{\partial^2\mathbf{r}}{\partial\theta\partial\phi}\cdot\mathbf{n} = 0 \qquad \text{(axisymmetric case)}$$

$$N = \frac{\partial^2\mathbf{r}}{\partial\phi^2}\cdot\mathbf{n} = R\sin(\theta)\left(\cos(\theta)u_\theta - \sin(\theta)\left(1+u-\frac{1}{2}u_\theta^2\right)\right) \tag{S11}$$

and total curvature $J$ is

$$J = -\frac{GL-2FM+EN}{EG-F^2} \tag{S12}$$

here with the sign convention flipped such that the vesicle has positive curvature with sense determined by a protein interacting with its outer leaflet. To second order $J$ is:

$$J = \frac{2}{R} - \frac{1}{R}\left(\cot(\theta)u_\theta + u_{\theta\theta} - 2u\left(-1+u+\cot(\theta)u_\theta+u_{\theta\theta}\right)\right) + \mathcal{O}[u^3] \tag{S13}$$

$$J = \frac{2}{R} - \frac{1}{R}\left(2u+\nabla^2u-2u^2-2u\nabla^2u\right) + \mathcal{O}[u^3] \tag{S14}$$

where we have used the expression for the constant-radius axisymmetric spherical Laplacian

$$\nabla^2u = \cot(\theta)u_\theta + u_{\theta\theta} \tag{S15}$$

The curvature-weighted area element $JdA$ simplifies to

$$JdA = R\left(2+\left(2u-\cot(\theta)u_\theta-u_{\theta\theta}+u_\theta^2\right)\right)\mathrm{d}\Omega \tag{S16}$$

$$= R\left(2+\left(2u-\nabla^2u+u_\theta^2\right)\right)\mathrm{d}\Omega \tag{S17}$$

The reduction to $\nabla^2u$ is critical as the spherical harmonics employed below for $u$ are eigenvalues of $\nabla^2$.

$$\nabla^2Y_l^0(\theta) = -l(l+1)Y_l^0(\theta) \tag{S18}$$

and as such integrals reduce cleanly, for example:

$$\int (\nabla^2 Y_l^0(\theta,\phi))^2 \mathrm{d}\Omega = (l\,(l+1))^2 \tag{S19}$$

where $Y_l^0(\theta)$ is normalized:

$$\int \left(Y_l^0(\theta)\right)^2 \mathrm{d}\Omega = 1. \tag{S20}$$

Spontaneous curvature $J_0$ is spread over an area $A_p$ at the polar cap. For a simple closed form expansion of the energy for weak perturbations $RJ_0$, we retain terms only to linear order in $u$. Integration of the energy $H$,

$$H = \oint_{\mathrm{cap}} \frac{\kappa}{2}\left(J - J_0\right)^2 \mathrm{d}A + \int_{\mathrm{vesicle}} \frac{\kappa}{2}\left(J\right)^2 \mathrm{d}A - \oint_{\mathrm{cap}} \frac{\kappa}{2}\left(J\right)^2 \mathrm{d}A \tag{S21}$$

is then split between the cap and the bulk with the term

$$\oint_{\mathrm{cap}} \kappa J J_0 \mathrm{d}A \tag{S22}$$

determining the force acting as a result of cap spontaneous curvature $J_0$. Integrals are computed over the cap region from $\theta = 0$ to $\theta = \theta_0$ with

$$x_0 = \cos(\theta_0) \tag{S23}$$

$$\oint_{\mathrm{cap}} \kappa J J_0 \mathrm{d}A = R\kappa J_0 \oint_{\mathrm{cap}} \left(2 + (2u - \nabla^2 u)\right) \mathrm{d}\Omega + \mathcal{O}[u^2] \tag{S24}$$

$$\oint_{\mathrm{cap}} \kappa J J_0 \mathrm{d}A = \underbrace{4\pi R\kappa J_0\left(1 - x_0\right)}_{u \text{ independent}} + \Lambda a_l \tag{S25}$$

(S26)

with

$$u\left(\theta,\phi\right) = \sum_l a_l Y_l^0(\theta,\phi) \tag{S27}$$

and

$$\Lambda = R\kappa J_0 \left( \underbrace{2\sqrt{\frac{\pi}{2l+1}}\left(P_{l-1}(x_0) - P_{l+1}(x_0)\right)}_{2u \text{ term}} + \underbrace{(2l+1)l\sqrt{\frac{\pi}{2l+1}}\left(P_{l-1}\left(x_0\right) - x_0 P_l\left(x_0\right)\right)}_{-\nabla^2 u \text{ term}} \right) \tag{S28}$$

where here $P_l(x_0)$ is the Legendre polynomial. At first order in $u$ it is not necessary to consider the area-weighted $J_0^2$ region of the cap, as the response of the cap-region area does not affect the deformation amplitude $u$ at first order.

The total vesicle elastic constant is the well-known value

$$K = \kappa(l-1)(l+2)l(l+1) \tag{S29}$$

with the energy for mode $l$

$$E_l = \frac{1}{2}K a_l^2 - \Lambda a_l \tag{S30}$$

such that the linear response of the harmonic amplitude $a_l$ is determined to be

$$a_l = \frac{\Lambda}{K} \tag{S31}$$

which reduces to

$$a_l = \frac{RJ_0\sqrt{\pi(1-x_0^2)(2l+1)}(2+l+l^2)}{l^2(1+l)^2(-2+l+l^2)} P_l^1(x_0) \tag{S32}$$

where $P_l^1(x_0)$ is the associated Legendre polynomial and modes $l$ are independent at first order in $u$.

Given the effect of the cap spontaneous curvature, the *change* in curvature (not including the curvature of the resting vesicle) for a particular spherical harmonic $l$ is estimated by integrating over its first region of same-sign curvature, equivalent to setting the limit of integration on $\theta$ to the first root of the Legendre polynomial, here is estimated by Bruns' formula to be $\frac{3\pi}{4l+2}$.

$$\Delta\bar{J}_l = \frac{1}{2\pi R^2(1-x_0')} \oint J \mathrm{d}A \tag{S33}$$

where $4\pi R^2(1-x_0')$ is the area of the integration region and $J$ is computed applying Eq. S17 to first order in $u$ with $a_l$ set as in Eq. S32. Note here that $x_0 = \cos(\theta_0)$ defines the region of the cap with spontaneous curvature $J_0$ and $x_0' = \cos(\theta_0')$ defines the integration region over the vesicle.

$$\Delta \bar{J}_l = \frac{a_l}{2\pi R^2(1-x_0'')} \left( \underbrace{R\, 2\sqrt{\frac{\pi}{2l+1}} \left(P_{l-1}(x_0') - P_{l+1}(x_0')\right)}_{2u \text{ term}} + \underbrace{R\,(2l+1)l\sqrt{\frac{\pi}{2l+1}} \left(P_{l-1}\left(x_0'\right) - x_0' P_l\left(x_0'\right)\right)}_{-\nabla^2 u \text{ term}} \right) \tag{S34}$$

or

$$\Delta \bar{J}_l = \frac{1}{2\pi R^2(1-x_0')} J_0 \frac{(1+2l)(2+l+l^2)^2}{l^3(1+l)^3(-2+l+l^2)} \pi R^2 \sin(\theta_0)\sin(\theta_0') P_l^1(\cos(\theta_0)) P_l^1(\cos(\theta_0')) \tag{S35}$$

again with

$$\theta_0' = \frac{3\pi}{4l+2} \tag{S36}$$

If the patch area $A_p$ is assumed to be small compared to the vesicle size then

$$\sin(\theta_0) P_l^1(\cos(\theta_0)) \approx \frac{-l(l+1)A_p}{2\pi R^2} \tag{S37}$$

yielding for $l = 2$

$$\Delta \bar{J}_2 \approx \frac{0.495 A_p J_0}{R^2} \tag{S38}$$

where

$$A_p J_0 = \frac{1}{2} A_{\text{sim}} \langle J \rangle \tag{S39}$$

as reported in the main text for curvature sampling by the peptides. For an object of radius 50 nm, the expected curvature sampling for $\langle J \rangle A$=10 Å is $10^{-5}$ Å$^{-1}$.

---


[1] N. Michaud-Agrawal, E. J. Denning, T. B. Woolf, and O. Beckstein, Journal of Computational Chemistry **32**, 2319 (2011), _eprint: https://onlinelibrary.wiley.com/doi/pdf/10.1002/jcc.21787.

[2] R. J. Gowers, M. Linke, J. Barnoud, T. J. E. Reddy, M. N. Melo, S. L. Seyler, J. Domański, D. L. Dotson, S. Buchoux, I. M. Kenney, and O. Beckstein, SciPy 2016 (2016), 10.25080/Majora-629e541a-00e.

[3] A. J. Sodt, A. H. Beaven, O. S. Andersen, W. Im, and R. W. Pastor, Biophysical Journal **112**, 1198 (2017).

[4] W. Humphrey, A. Dalke, and K. Schulten, Journal of Molecular Graphics **14**, 33 (1996).

[5] Y. Sato, S. Nakajima, N. Shiraga, H. Atsumi, S. Yoshida, T. Koller, G. Gerig, and R. Kikinis, Medical Image Analysis **2**, 143 (1998).

[6] C. R. Harris, K. J. Millman, S. J. van der Walt, R. Gommers, P. Virtanen, D. Cournapeau, E. Wieser, J. Taylor, S. Berg, N. J. Smith, R. Kern, M. Picus, S. Hoyer, M. H. van Kerkwijk, M. Brett, A. Haldane, J. F. del Río, M. Wiebe, P. Peterson, P. Gérard-Marchant, K. Sheppard, T. Reddy, W. Weckesser, H. Abbasi, C. Gohlke, and T. E. Oliphant, Nature **585**, 357 (2020).

[7] P. Virtanen, R. Gommers, T. E. Oliphant, M. Haberland, T. Reddy, D. Cournapeau, E. Burovski, P. Peterson, W. Weckesser, J. Bright, S. J. van der Walt, M. Brett, J. Wilson, K. J. Millman, N. Mayorov, A. R. J. Nelson, E. Jones, R. Kern, E. Larson, C. J. Carey, I. Polat, Y. Feng, E. W. Moore, J. VanderPlas, D. Laxalde, J. Perktold, R. Cimrman, I. Henriksen, E. A. Quintero, C. R. Harris, A. M. Archibald, A. H. Ribeiro, F. Pedregosa, and P. van Mulbregt, Nature Methods **17**, 261 (2020).

[8] S. van der Walt, J. L. Schönberger, J. Nunez-Iglesias, F. Boulogne, J. D. Warner, N. Yager, E. Gouillart, and T. Yu, PeerJ **2**, e453 (2014).

[9] J. D. Hunter, Computing in Science & Engineering **9**, 90 (2007).
